# Telomeric SUMO level influences the choices of APB formation pathways and ALT efficiency

**DOI:** 10.1101/2025.01.16.633463

**Authors:** Rongwei Zhao, Allison Wivagg, Rachel M. Lackner, Jayme Salsman, Graham Dellaire, Michael J Matunis, David M. Chenoweth, Xiaolan Zhao, Huaiying Zhang

**Affiliations:** Department of Biology, Carnegie Mellon University, Pittsburgh, PA 15213, USA; Department of Chemistry, University of Pennsylvania, Philadelphia, PA 19014, USA; Department of Pathology, Dalhousie University, Halifax, Nova Scotia, B3H 4R2, Canada; Department of Biochemistry and Molecular Biology, Johns Hopkins University, Baltimore, MD 21205, USA; Molecular Biology Program, Memorial Sloan Kettering Cancer Center, New York, NY 10065, USA

## Abstract

Many cancers use an alternative lengthening of telomeres (ALT) pathway for telomere maintenance. ALT telomeric DNA synthesis occurs in ALT telomere-associated PML bodies (APBs). However, the mechanisms by which APBs form are not well understood. Here, we monitored the formation of APBs with time-lapse imaging employing CRISPR knock-in to track the promyelocytic leukemia (PML) protein at endogenous levels. We found APBs form via two pathways: telomeres recruit PML proteins to nucleate PML bodies de novo, or telomeres fuse with preformed PML bodies. Both nucleation and fusion of APBs require interactions between SUMO and SUMO interaction motifs (SIMs). Moreover, APB nucleation is associated with higher levels of SUMOs and SUMO-mediated recruitment of DNA helicase BLM, resulting in more robust telomeric DNA synthesis. Finally, further boosting SUMO levels at telomeres enhances APB nucleation, BLM enrichment, and telomeric DNA synthesis. Thus, high SUMO levels at telomeres promote APB formation via nucleation, resulting in stronger ALT activity.

## Introduction

Cancer cells need to overcome replication-associated telomere shortening and senescence to maintain proliferation during oncogenic stress (Okamoto & Seimiya, 2019). While most cancer cells achieve this via re-activating telomerase (Shay, 2016), ∼15% of the cancer cells utilize alternative lengthening of telomeres (ALT) through a recombination-based repair mechanism called break-induced replication (BIR) (Pickett & Reddel, 2015; J. M. Zhang & Zou, 2020; Dilley et al., 2016). ALT takes place in specialized PML nuclear bodies (PML NBs) called APBs (ALT telomere-associated PML NBs) that enrich both clustered telomeres and BIR proteins, thus serving as a hub for BIR-mediated telomere recombination (Brouwer et al., 2009; Chung et al., 2011; J. M. Zhang et al., 2019; Draskovic et al., 2009). The presence of APBs has provided a main strategy for ALT diagnosis, and dampening APB formation can potentially halt ALT cancer proliferation (Heaphy et al., 2011).

SUMOylation, the process by which the small ubiquitin modifier (SUMO) is conjugated to target proteins, has been shown to play a key role in APB formation and ALT (Osterwald et al. 2015; Min et al. 2019). Indeed, SUMOylation inhibitors can reduce APB levels and ALT cell growth, indicating their potential therapeutic effects against ALT cancers (Zhao et al., 2024). SUMOylation affects three groups of proteins important to APB formation and ALT. First, SUMOylation of telomeric proteins, such as TRF1, TRF2, TIN2, and RAP1, is required for APB formation (Brouwer et al., 2009; Chung et al., 2011; Potts & Yu, 2007). Second, SUMOylation of DNA repair factors, such as Rad52 and BLM, enables their recruitment to APBs (J.-M. Zhang et al., 2021). Third, SUMOylation of the PML protein, the scaffold component of the PML NBs, is critical for PML NBs and APB formation (Chung et al., 2011; Duprez et al., 1999; Tatham et al., 2008; Zhao et al., 2024). A prominent role of SUMOylation is to promote interactions amongst telomeric proteins, repair proteins, and the PML protein via interactions between SUMOs and SUMO interaction motifs (SIMs). Numerous SIMs present on these proteins and their SUMOylation at many sites permit multi-valent SUMO-SIM interactions, which can drive phase separation to form liquid condensates (Keiten-Schmitz et al., 2021; Raman et al., 2013; Shen et al., 2006; Zhang et al., 2020). Indeed, the importance of SIMs in APB functions is demonstrated as SIMs of DNA repair factors such as Rad51AP1 and BLM are required for their APB localization (Loe et al., 2020; Min et al., 2019). In addition, the SIM in the PML protein is also important for PML NBs formation and APB formation (Shen et al., 2006; Lallemand-Breitenbach and Thé, 2010; J.-M. Zhang et al., 2021).

While the requirement of SUMOylation for APB formation is well-established, important details are lacking regarding APB formation pathways and the roles of SUMOylation in this process. It is unclear whether APBs form via de novo nucleation of PML NBs on telomeres or via recruiting telomeres to pre-existing PML NBs. In addition, it is unknown how different SUMO paralogs and SUMO E3 ligases affect different APB formation pathways and if they do so by controlling SUMOylation of telomere-binding proteins, the DNA repair factors, or the PML protein. Addressing these questions requires advanced techniques allowing for live cell time-lapse imaging of APBs in a condition wherein new APB formation can be induced and monitored in real-time. In addition, the time-lapse imaging must be done at the endogenous PML protein level, as overexpressing the PML protein is known to affect PML NB formation (Corpet et al., 2020). One method that allows inducible APB formation is the TRF1-FokI system (Cho et al., 2014; Dilley et al., 2016). In this system, FokI is fused with TRF1 (telomere repeat factor 1), DD (destabilization domain), and ER (modified estrogen receptor) domain. The addition of 4-hydroxytamoxifen (4-OHT) and shield1 induces FokI nuclease release to nucleoplasm and translocation to telomeres (Fig. 1B). Localization of FokI-TRF1 to telomeres then triggers DNA breaks exclusively at telomeres, leading to the formation of APBs with all endogenous APB characteristics, such as telomere clustering and telomeric DNA synthesis. Another method for APB formation with temporal control entails tethering SUMO to telomeres using the chemical-induced protein-dimerization system (Fig. 1H) (Lackner et al., 2022; Zhao et al., 2021). In this system, the dimerizer TMP (trimethoprim)-Fluorobenzamide-Halo ligand (TFH) is composed of chemically linked TMP and Halo ligand that can interact with the *E. coli* dihydrofolate reductase (eDHRF) and the Halo-enzyme, respectively. Adding the dimerizer to cells expressing SUMO/SIM fused to eDHRF and TRF1 fused to Halo-enzyme will target SUMO/SIM to telomeres to form APBs on demand. This enables live-cell time-lapse imaging and prevents the toxicity caused by constitutive telomere targeting. Importantly, with CRISPR technology, we engineered an ALT-positive cell line U2OS where the endogenous PML protein is tagged by CRISPR knock-in with a cDNA encoding the fluorescent protein Clover (Pinder et al., 2015), allowing us to follow APB formation without overexpressing the PML protein.

**Fig. 1.**
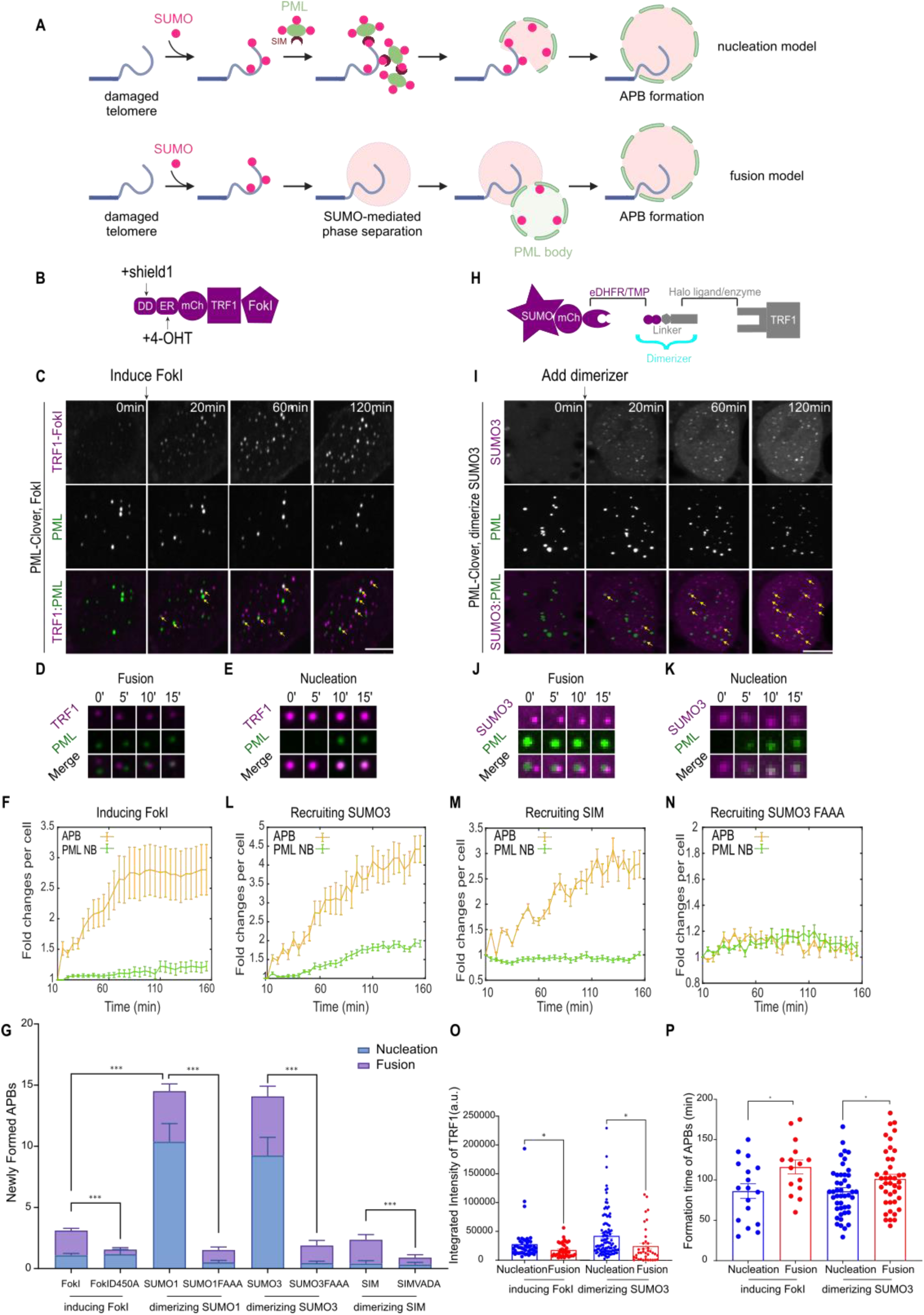
APBs are formed by both fusion and nucleation. **(A)** Schematic for APB nucleation and fusion model. **(B)** Schematic for mCherry-TRF1-FokI. FokI was fused with TRF1, DD (destabilization domain) and ER (modified estrogen receptor) domain. By adding 4-Hydroxyestradiol (4-OHT, a selective estrogen receptor modulator that binds to ER) and shield1 (stabilizes protein tagged with the DD domain), FokI will be released to nucleoplasm and localized to telomeres. **(C)** Representative images of cells with endogenous PML protein labeled with Clover and expressing mCh-TRF1-FokI with the treatment of 4-OHT to induce DNA damage at indicated time points. Yellow arrows indicate APBs. **(D, E)** Images with APBs formed by fusion and nucleation after FokI induction, 5 min between images. **(F)** Quantification of APB and PML NB numbers per cell after FokI induction. **(G)** Quantification of newly formed APBs per cell after inducing FokI or dimerizing SUMO/SIM for 2 hours. More than 30 cells were analyzed in each group from three independent experiments. **(H)** Dimerization schematic: SUMO or SIM was fused to mCherry and eDHFR, and TRF1 was fused to 3xHalo enzyme. The dimerizer, TMP (trimethoprim)-Fluorobenzamide-Halo ligand (TFH), can interact with eDHFR and the Halo enzyme to dimerize SUMO or SIM to TRF1. TRF1 was not fluorescently labeled, and SUMO3 was used to indicate telomeres in the images. **(I)** Representative images of cells with endogenous PML labeled with Clover, expressing mCh-eDHFR-SUMO3 and 3xHalo-TRF1 with the addition of the dimerizer at indicated time points. Yellow arrows indicate APBs. **(J, K)** Images with APBs formed by fusion and nucleation after dimerizing SUMO3 to TRF1, 5 min between images. **(L)** Quantification of APB and PML NB numbers per cell after dimerizing SUMO3. **(M)** Quantification of APB and PML NB numbers per cell after dimerizing SIM. **(N)** Quantification of APB and PML NB numbers per cell after dimerizing SUMO3(FAAA). **(O)** The integrated intensity of TRF1-FokI and TRF1-SUMO3 comparing nucleation and fusion after inducing FokI or dimerizing SUMO3 for 2 hours. Each dot represents one cell in three independent experiments, more than 42 cells were analyzed in each group. **(P)** APB formation time for nucleation and fusion after inducing FokI or dimerizing SUMO3 for 2 hours. Each dot represents one cell in three independent experiments, more than 40 cells were analyzed in each group. Scale bars, 5 μm or 1 μm for the cropped images. Numbers in (F)(L)(M)(N) were normalized by the number at the first time point for each cell, more than 16 cells per group, in three independent experiments.

Taking advantage of the TRF1-FokI system, the SUMO/SIM telomere targeting method, and the PML-Clover U2OS cell line, we monitored APB formation dynamics using live cell time-lapse imaging at endogenous PML protein level. We examined two possible pathways for APB formation. First, given that the PML protein is decorated with SIMs and conjugated with SUMO moieties (Kamitani et al., 1998; Shen et al., 2006), we envision it can be recruited to telomeres that are enriched with SUMO/SIM via SUMO-SIM interactions to form PML NBs on telomeres, and thus APBs. This model of de novo formation of APBs via nucleating PML NBs on telomeres is referred to as the APB nucleation model (Fig. 1A top). Alternatively, considering that SUMO mediates phase separation of DNA repair factor located on ALT telomeres without PML (Zhao et al., 2024), it is possible that thus-formed condensates can fuse with pre-existing PML NBs, which are also condensates enriched with SUMO, to form APBs. We refer to this hypothesis entailing two types of condensates coalescing as the APB fusion model (Fig. 1A bottom). We provide evidence that both mechanisms are present in the same cell and both require SUMO-SIM interactions. We detail the contributions of SUMO/SIM from the PML protein, different SUMO paralogs, and SUMO E3 ligases for both APB formation and ALT telomeric DNA synthesis. Importantly, we report a set of observations indicating that SUMO levels at telomeres influence the two APB formation mechanisms. First, we find that APB nucleation occurs more often on telomeres with higher SUMO levels compared with APB fusion. Second, boosting the telomeric SUMO level can promote more APB nucleation. Third, APB nucleation is more sensitive to reducing telomere SUMO levels. These data are consistent with each other and indicate that SUMO level is a determining factor in the choices of APB formation mechanisms. Finally, we show the functional difference of these two APB formation mechanisms, with APB nucleation leading to more robust telomere synthesis. Collectively, our data provides direct visualization of two APB formation routes, reveals differential ALT potentials of these two pathways, and uncovers how the two APB formation choices depend on SUMO levels at telomeres.

## Results

### Telomere breaks induce APB formation via both nucleation and fusion

We used the TRF1-FokI system to induce APB formation in the PML-Clover U2OS cell line and followed the dynamics of newly formed APBs using time-lapse imaging. Since PML is the major structural component of PML NBs, PML-Clover-enriched foci represent PML NBs (Pinder et al., 2015). Accordingly, PML-Clover foci that are enriched in mCherry-TRF1-Fok1 represent APBs. Live imaging showed an increase in the number of APBs over time, indicating that new APBs were formed after inducing FokI (Fig. 1C, Movie 1). As a control, inducing the enzymatically dead mutant of FokI (D450A) failed to increase APB numbers (Fig. 1G, Fig. S1A, B), confirming that new APB formation was induced by telomeric-specific DNA breaks as seen previously (Cho et al., 2014).

Tracing new APB formation after telomere break induction revealed both fusion and nucleation events. We define fusion events as the collision of FokI-TRF1 marked telomere foci with the pre-existing PML foci, which resulted in the sudden appearance of APBs enriched in both PML and TRF1 at telomeres (Fig. 1D, Movie 1). We define nucleation events as a gradual gain of PML signals on telomeres without a collision of pre-existing PML foci with telomere foci (Fig. 1E, Movie 1). To confirm the visual inspection, we quantified the change in the total number of PML NBs over time. We expect the PML NB number to remain unchanged for APB formation via fusion because it is simply a re-localization of existing PML NBs to telomeres. On the other hand, we anticipate the PML NB number to increase for APB formation via nucleation because new PML NBs are formed on telomeres. Confirming that APBs are formed by nucleation after TRF1-FokI induction, we observed an increase in the PML NB number (Fig. 1F). This was not seen when using TRF1-FokI (D450A) construct as expected (Fig. S1B).

Quantifying the number of APBs formed via fusion versus nucleation revealed that fusion events were about twice as frequent compared to nucleation events during the 2 hours following FokI induction (Fig. 1G). Overall, telomeres that nucleated APBs had a higher intensity than those engaged in APB formation via fusion (Fig. 1O). Quantifying the time of APB formation revealed that nucleation of APBs happened faster than fusion (Fig. 1P). While the underlying reason for this difference is unclear, a simple interpretation is that PML proteins diffuse faster than PML NBs in the nucleoplasm. These results show that APBs can form via nucleation and fusion in the same cell, but these pathways differ in terms of speed and frequency as well as the level of telomere signals. We conclude that in the context of telomere breaks, APB fusion occurs more frequently, while APB nucleation happens faster and is favored by stronger telomeric signals.

### Targeting SUMO to telomeres potently induces APB nucleation and fusion

To understand the dependence of the two APB formation pathways on SUMO, we used the SUMO-telomere targeting system that can induce new APBs in the PML-Clover cells (Fig. 1H). Here, we targeted SUMO3, one of the three human SUMO paralogs, and used its non-conjugatable form (C-terminal di-Gly motif mutated to di-Val) to prevent conjugation (Banani et al., 2016). We found that recruiting SUMO3 to telomeres resulted in APB formation via both nucleation and fusion, as shown by both live imaging data and quantifications of APB and PML NB numbers over time (Fig. 1I-L, Movie 2).

As seen in the TRF1-Fok1 system, APB nucleation happened faster than fusion in the SUMO-telomere tethering system (Fig. 1P). The SUMO targeting systems further revealed that APB nucleation tended to occur on telomeres with stronger SUMO signals (Fig. 1O). Considering that stronger telomere signals in U2OS cells are more enriched with SUMO (Zhao et al., 2024), this finding is consistent with the observations made in the TRF1-FokI system, wherein APB nucleation tended to occur on brighter telomeres (Fig. 1O).

Strikingly, new APBs generated via both pathways upon SUMO tethering to telomeres were more frequent than those seen after TRF1-FokI induction, indicating that SUMO-telomere targeting is more potent in inducing APBs (Fig. 1G). This better potency is related to the observation that enrichment of SUMO is a downstream event after telomere break induction in ALT (H. Zhang et al., 2020). Consequently, a direct increase in telomeric SUMO level can be more effective in inducing APBs than telomere breaks. Given that higher SUMO levels at telomeres favor APB nucleation (Fig. 1G), it is not surprising that most APBs formed upon targeting SUMO to telomeres were via nucleation, where both PML NB and APB number increased after dimerizing SUMO to telomeres (Fig. 1G, L).

Collectively, these data indicate that both telomere breaks and SUMO-telomere tethering led to new APB formation via fusion and nucleation, with nucleation being the faster event. The observed differences between the two systems can be unified by the theory that APB nucleation is favored at telomeres with higher SUMO levels. We note that SUMO level rather than the identity of SUMO paralogs is more important in determining APB formation via nucleation or fusion, as targeting SUMO1 to telomeres led to similar outcomes as those described for SUMO3 targeting (Fig. 1G).

### Both pathways of APB formation require SUMO-SIM interactions

As both APB nucleation and fusion can occur upon SUMO tethering at telomeres, we asked whether the role of SUMO in these processes requires its binding to SIM. To this end, we recruited SUMO1 or SUMO3 mutants that could not interact with SIM (FKVK in SUMO1 and FKIK in SUMO3 mutated to FAAA) to telomeres (Banani et al., 2016). We found reduced APB formation via both nucleation and fusion compared with their wild-type counterparts (Fig. 1G, N, Fig. S1C, Movie 3). Nucleation decreased more dramatically than fusion for both mutants (Fig. 1G), indicating that nucleation of APBs relies more heavily on SUMO-SIM interactions at the telomere compared to fusion. A similar conclusion was reached when examining telomere targeting of SIM or its SUMO-binding mutant (VIDL mutated to VADA) (Banani et al., 2016), since recruiting SIM, but not its mutant, led to APB formation via nucleation and fusion (Fig. 1G, M, Fig. S1D-F). Collectively, these results support the notion that SUMO-SIM interactions are critical for both APB formation pathways.

In the above experiments, we noticed that SIM targeting was less potent in inducing APB nucleation than SUMO targeting, but both had similar effects on APB fusion (Fig. 1G, Fig. S1D, Movie 4). This was shown by the increase in APB numbers without changes in PML NB numbers after SIM recruitment (Fig. 1M). One interpretation of these results is that PML protein recruitment to telomeres during APB nucleation relies more on interaction between PML and SUMO at telomeres, while fusion between PML NBs and telomeres does not have this preference (Fig. 1G, Fig. S1E, F).

### Both SUMOylation sites and SIMs of the PML protein are important for APB formation

To further understand the role of SUMO and SIM in APB formation, we focused on the PML protein, which contains multiple SUMOylation sites and one SIM (Kamitani et al., 1998; Shen et al., 2006). This choice is based on the consideration that the PML protein is key for the PML NB and APB formation. We expressed PML-IV, one isoform of the PML protein, and its variants, which are mutated at their SUMOylation or SIM sites, in PML knockout (PML KO) U2OS cells. The SUMOylation mutant, PML(SUMO-), was generated by mutating its SUMOylation sites K65, K160, and K442 to arginine, and the SIM mutant, PML(SIM-), was generated with the SIM site deleted (Banani et al., 2016; Shen et al., 2006). We found that PML-IV rescued APB fusion and nucleation in PML KO cells after SUMO and SIM tethering (Fig. 2B). However, PML(SUMO-) and PML(SIM-) only showed a low level of rescue of APB formation via nucleation and fusion (Fig. 2A, B, Fig. S2B, D, E), indicating that both pathways of APB formation require the SUMOylation sites and SIM on the PML protein. We noticed that the PML(SUMO-) was more defective than PML(SIM-) in this assay (Fig. 2B, Fig. S2A, C, F, G). One interpretation for this difference is that PML SUMOylation is more important than its SIM for APB formation.

**Fig. 2.**
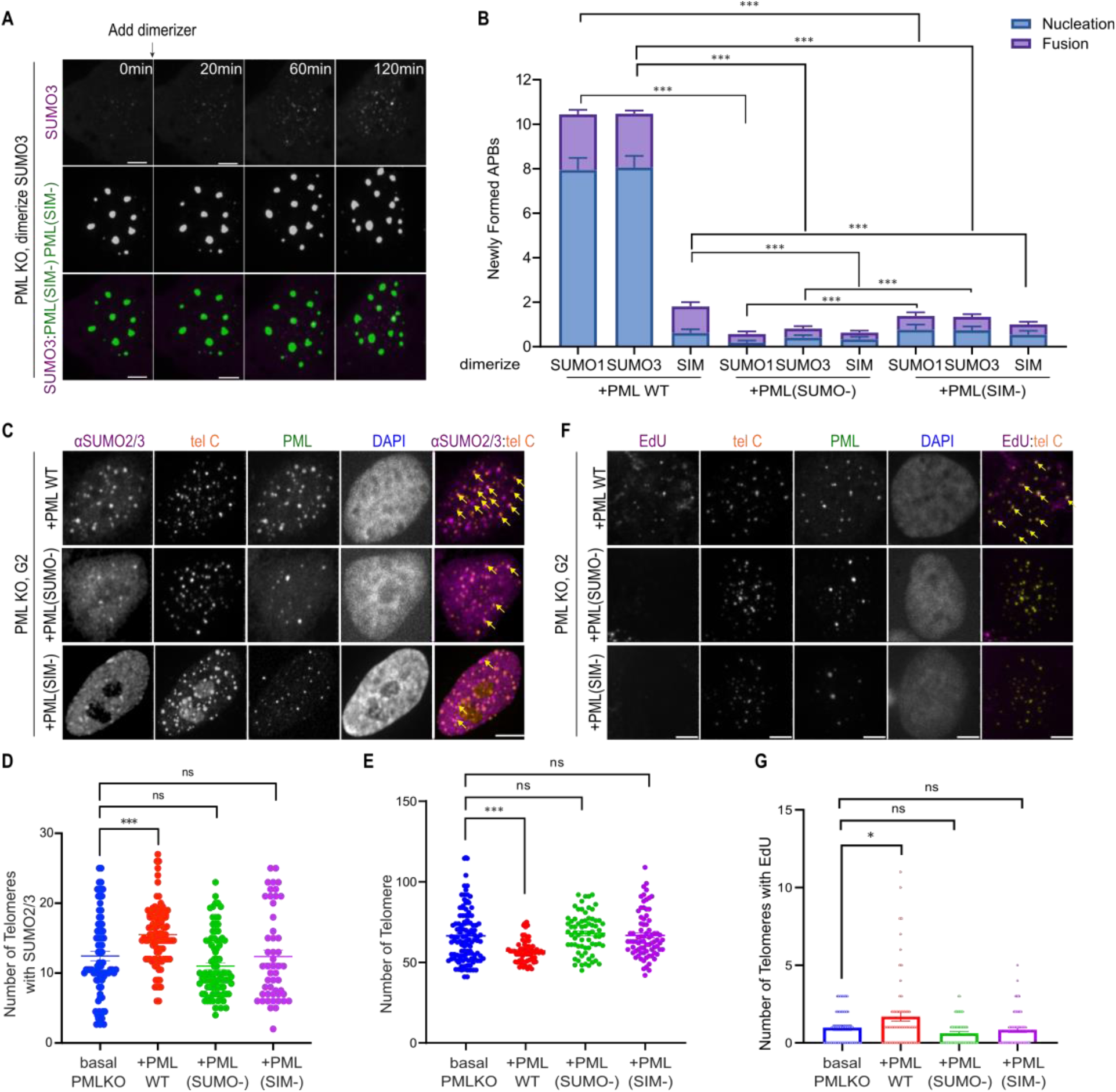
PML protein contributes to ALT via its SUMOylation sites and SIM. **(A)** Representative images of PML KO cells expressing mCh-eDHFR-SUMO3, 3xHalo-TRF1 and PML(SIM-) after adding the dimerizer to dimerize at indicated time points. **(B)** Quantification of newly formed APBs per cell after recruiting SUMO/SIM expressing PML(SUMO-) or PML(SIM-) in PML KO cells. More than 23 cells were analyzed in each group from three independent experiments. **(C)** Representative images and **(D)** quantification of SUMO2/3 localization on telomeres in G2-arrested PML KO after overexpressing PML WT, PML(SUMO-) or PML(SIM-). Each dot represents one cell in three independent experiments, more than 52 cells were analyzed in each group. Yellow arrows indicate SUMO2/3 localized on telomeres. **(E)** Quantification of telomere number in G2 arrested PML KO after overexpressing PML WT, PML(SUMO-) or PML(SIM-). Each dot represents one cell in three independent experiments, more than 41 cells were analyzed in each group. **(F)** Representative images and **(G)** quantification of EdU signal on telomeres in G2 arrested PML KO after overexpressing PML WT, PML(SUMO-) or PML(SIM-). Each dot represents one cell in three independent experiments, more than 44 cells were analyzed in each group. Yellow arrows indicate EdU signal on telomeres. Scale bars, 5 μm.

These results led us to examine the ability of PML-IV and its SUMOylation and SIM mutants to rescue ALT defects in PML KO cells at the G2 phase where ALT is active. We assessed key ALT features, including SUMO1 and SUMO2/3 levels at telomeres (Fig. 2C, D, Fig. S2H, I), APB formation (Fig. S2J), telomere clustering (inversely correlated with telomere numbers) (Fig. 2E), and telomeric DNA synthesis (pulsed with fluorescent EdU, 5-Ethynyl-2-deoxyuridine) (Fig. 2F, G). As expected, PML-IV conferred rescue of all defects examined. However, PML(SUMO-) and PML(SIM-) did not significantly rescue any of the defects. These results support the previous studies showing that both SUMOylation sites and the SIM on the PML protein are important for APB formation and that SUMOylation sites are required for c-circle generation, another phenotype of ALT cells (J.-M. Zhang et al., 2021; Loe et al., 2020). In addition, our data also showed that PML(SUMO-) and PML(SIM-) exhibited equal importance for ALT phenotypes, except that the former was more defective in APB formation (Fig. 2B, Fig. S2J). Combining the above results, we conclude that both SUMOylation and SIM on the PML protein are important for ALT features, and SUMOylation sites have a stronger role in APB formation.

### SUMO2/3 makes more prominent contributions to APB formation and ALT than SUMO1

While targeting SUMO1 and SUMO3 to telomeres can induce APB formation equally, it is unclear if the endogenous SUMO paralogs have differential effects on APB formation and ALT. To address this question, we used siRNA to knock down SUMO1 and SUMO2/3 (targeted by the same siRNA) in U2OS cells (Fig. S3A-C). SUMO1 and SUMO2/3 knockdown efficiency were confirmed using western blot (Fig. 3A) and immunofluorescence (Fig. S3A-C). Consistent with previous findings, both knockdowns reduced PML NB numbers (Fig. 3B, C). In addition, both reduced ALT features in G2-arrested U2OS cells, including APB number (Fig. 3B, C), telomere clustering (Fig. S3D), and telomeric DNA synthesis (Fig. 3B, D), with SUMO2/3 knock down exhibiting stronger effects than SUMO1 knockdown (Fig. 3B-E, Fig. S3D). We used SUMO1 knock-out (SUMO1 KO), SUMO2 knock-out (SUMO2 KO), and SUMO2 knock-out with SUMO2 rescue (SUMO2 KO+SUMO2) U2OS cell lines to confirm the observations and to conduct rescue experiments (Wang et al., 2023). The SUMO1 and SUMO2 knock-out were confirmed by immunofluorescence (Fig. S4A-D) and western blot (Fig. S4M). Similar to siRNA knockdown, there were fewer APBs (Fig. S4E, G), fewer PML NBs (Fig. S4F), less enrichment of SUMO/SIM containing DNA repair factors including PCNA and RPA (Fig. S4J-L), less telomere clustering (Fig. S4H), and less telomeric DNA synthesis (Fig. S4I) in SUMO1 and SUMO2 KO cells than the SUMO1 and SUMO2 containing control U2OS cells. In addition, the defects were rescued in SUMO2 KO+SUMO2 cells (Fig. S4C-L). In all assays, SUMO2 KO caused more severe phenotypes than SUMO1 KO (Fig. S4E, G, H, I), consistent with the siRNA results and showing that SUMO2 has a more prominent role in APB formation and ALT than SUMO1 (Fig. 3B-D). Furthermore, similar findings were also observed in SUMO1 and SUMO2/3 siRNA knockdown experiments in U2OS cells lacking PML (Fig. 3D, E, Fig. S3B-D). We therefore conclude that SUMO2/3 contributes to ALT more than SUMO1 in both PML-dependent and -independent pathways.

**Fig. 3.**
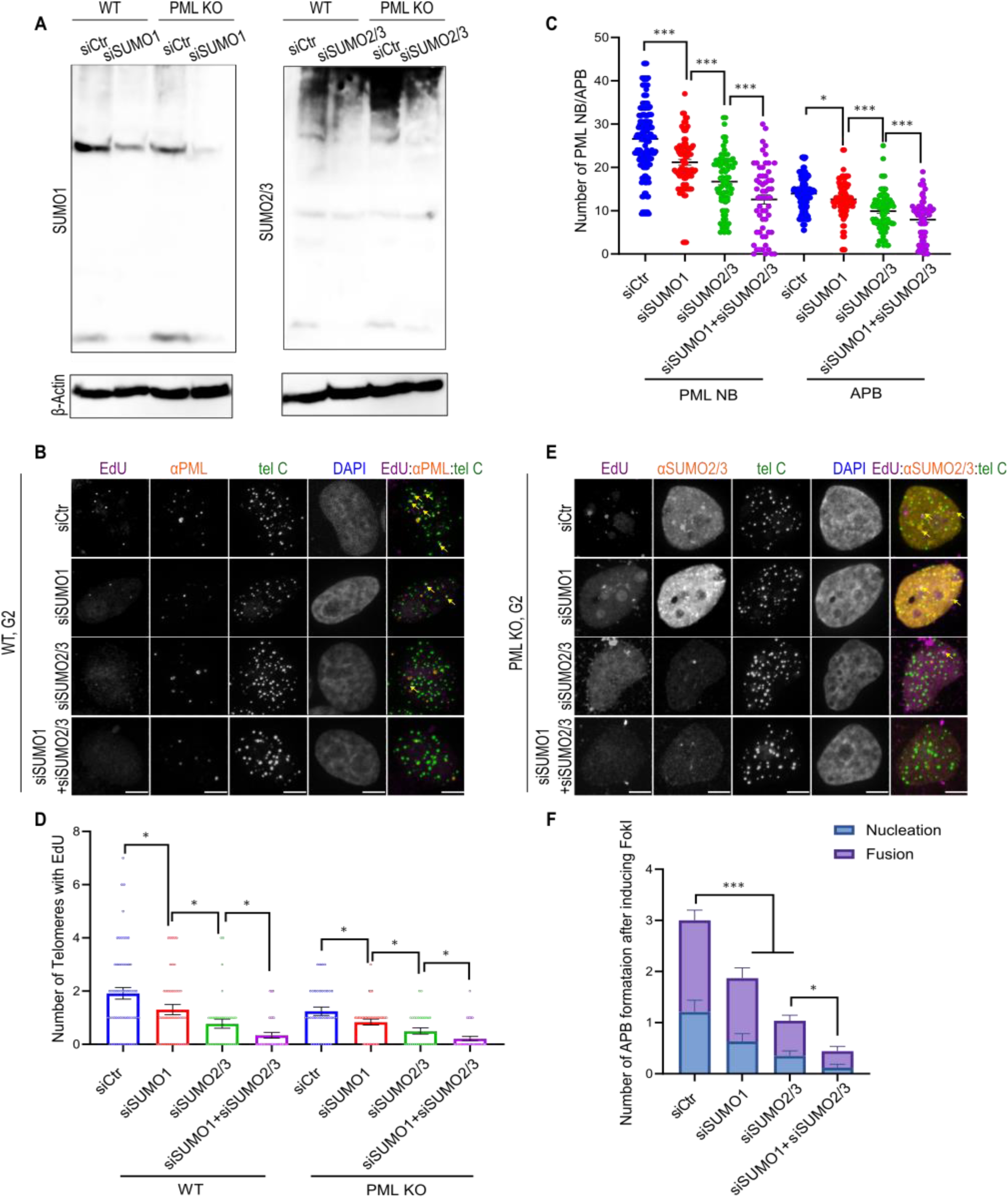
SUMO2/3 contributes to APB formation and ALT phenotypes more than SUMO1. **(A)** Western blot of SUMO1, SUMO2/3 (same antibody), or β-Actin after transfection of control siRNA, siSUMO1 or siSUMO2/3 in WT U2OS and PML KO U2OS cells for two days. **(B)** Representative images of telomeres with EdU, PML and **(E)** SUMO2/3 staining in G2-arrested U2OS WT cells and PML KO cells after transfection of control siRNA, siSUMO1 or siSUMO2/3 for two days. Yellow arrows indicate EdU signals at telomeres. **(C)** Quantification for the number of PML NBs and APBs in G2 arrested U2OS WT cells after transfection of control siRNA, siSUMO1 or siSUMO2/3 for two days. Each dot represents one cell in three independent experiments, more than 46 cells were analyzed in each group. **(D)** Quantification of the number of EdU foci on telomeres in G2 arrested U2OS WT and PML KO cells after transfection of control siRNA, siSUMO1 or siSUMO2/3 for two days. Each dot represents one cell in three independent experiments, more than 50 cells were analyzed in each group. **(F)** Quantification of newly formed APBs per cell after FokI induction for 2 hours, with transfection of control siRNA, siSUMO1 or siSUMO2/3 for 2 days. More than 23 cells were analyzed in each group from three independent experiments. Scale bars, 5 μm.

We found that, in PML-Clover-containing cells, knocking down SUMO1, SUMO2/3, or both greatly reduced APB nucleation and fusion after FokI induction (Fig. 3F), with a stronger effect seen for nucleation. This indicates that SUMO has more prominent roles in APB nucleation than fusion, agreeing with the data presented above using the SUMO-telomere recruitment system (Fig. 1G). Again, SUMO2/3 loss reduced nucleation and fusion to a greater extent than SUMO1 loss (Fig. 3F), further supporting a more prominent role of SUMO2/3 in APB formation pathways. Taken together, data using different APB induction and genetic perturbation systems are consistent with each other in supporting the important roles of SUMO1 and SUMO2/3 in ALT phenotypes. The greater effects of SUMO2/3 than SUMO1 are likely because SUMO2/3, but not SUMO1, can form polySUMO chains, possibly leading to a more efficient increase in SUMO levels on telomeres.

### MMS21 contributes to SUMO enrichment on telomeres without PML more than PIAS4

Two SUMO E3 ligases, namely MMS21 and PIAS4 (Brouwer et al., 2009; Chung et al., 2011; Potts & Yu, 2007; J.-M. Zhang et al., 2021), have been implicated in ALT; however, whether they have differential effects on APB formation and ALT is unknown. To test this, we examined the consequences of their knockdown in U2OS cells. Western blot confirmed that both knockdowns were successfully achieved using siRNA (Fig. 4A). As expected, both knockdowns impaired ALT features in G2 cells, including SUMO1/2/3 localization on telomeres (Fig. 4B, C, Fig. S5D), APB number (Fig. S5A, B), telomere clustering (Fig. S5E), localization of SUMO/SIM containing DNA repair factors PCNA and RPA to telomeres (Fig. S5F-I), and telomeric DNA synthesis (Fig. 4B, D). Both also reduced PML NB numbers (Fig. S5C), indicating that the two ligases affect PML NBs beyond APBs. Observed defects were quantitatively similar between the two knockout treatments and the combined treatment had an additive effect (Fig. 4C, D, Fig. S5B-E). Similar observations were made in PML KO cells, except that compared with PIAS4 knockdown, MMS21 knockdown more severely impaired SUMO localization on telomeres (Fig. 4C, E, Fig. S5D) and telomere clustering (Fig. S5E). This indicates that among the three groups of SUMOylation substrates that are important for ALT function, namely telomere binding proteins, DNA repair factors, and the PML protein, MMS21 affects the SUMOylation of the former two groups more than PIAS4. Finally, monitoring PML-Clover upon TRF1-Fok1 induction showed that MMS21 and PIAS4 knockdown had comparable effects in reducing APB fusion and nucleation, with their combined knockdown showing an additive effect (Fig. 4F). Collectively, these results indicate that MMS21 and PIAS4 make separate yet comparable contributions to APB formation and ALT but MMS21 has a more prominent role in SUMO accumulation at telomeres without PML.

**Fig. 4.**
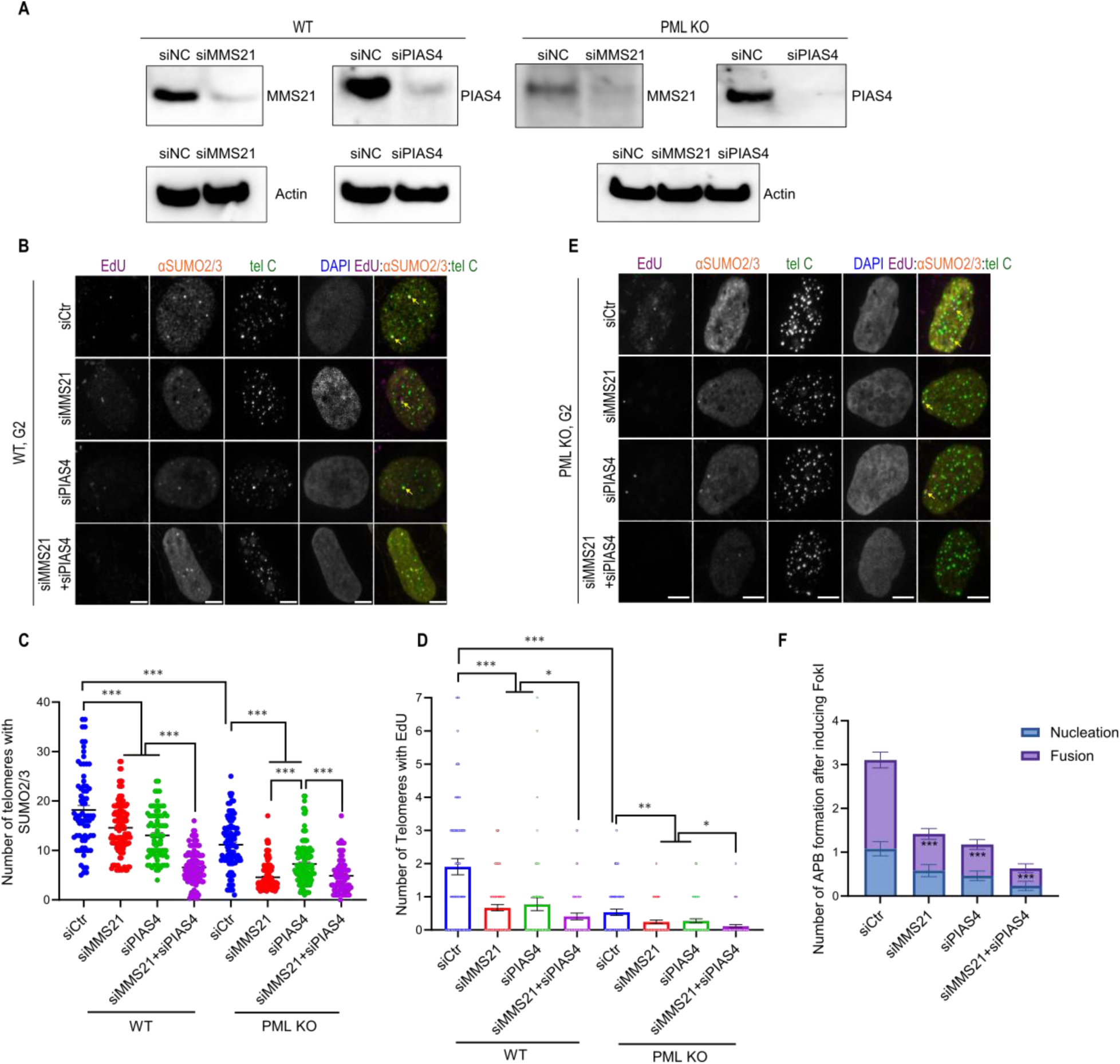
MMS21 contributes to SUMO level on telomeres more than PIAS4 in the absence of PML. **(A)** Western blot of MMS21, PIAS4 or β-Actin after transfection of control siRNA, siMMS21 or siPIAS4 in U2OS and PML KO cells for two days. **(B)** Representative images of telomeres with EdU, SUMO2/3 staining in G2 arrested U2OS WT, and **(E)** PML KO cells after transfection of control siRNA, siMMS21 or siPIAS4 for two days. Yellow arrows indicate EdU signals at telomeres. **(C)** Quantification of SUMO2/3 localization and **(D)** EdU signal on telomeres in G2 arrested U2OS WT and PML KO cells after transfection of control siRNA, siMMS21, or siPIAS4 for two days. Each dot represents one cell in three independent experiments, more than 46 cells were analyzed in each group. **(F)** Quantification of newly formed APBs per cell after FokI induction for 2 hours, with transfection of control siRNA, siMMS21, or siPIAS4 for 2 days. More than 20 cells were analyzed in each group from three independent experiments. Scale bars, 5 μm.

### Recruiting SUMO ligases to telomeres forms APBs mainly via fusion

To test the ligases’ differential roles in SUMO function at telomeres further, we used the protein-tethering system described above to target MMS21 or PIAS4 to telomeres. To assess active SUMOylation on telomeres, we co-expressed HA-SUMO3, pulled down HA, and blotted TRF2 after dimerizing MMS21 or PIAS4 to TRF1. We confirmed that recruitment of either ligase could increase SUMOylated TRF2, but not their catalytic dead variants, namely MMS21(C215A) and PIAS4(C342A) (Fig. S6I, J). Immunofluorescence also confirmed an increase in SUMO localization on telomeres (Fig. 5A, Fig. S6B-D), indicating there was elevated telomere SUMOylation, which was not seen with MMS21(C215A) and PIAS4(C342A) (Fig. 5A, Fig. S6B-D). Both tethering also led to increases in telomere clustering (Fig. S6F) and APB formation (Fig. 5C, D, Fig. S6E). Live imaging showed that both systems induced signatures of phase separation, including fusion events (Fig. S7A, D, Movie 5, 6), increased telomere intensity (Fig. S7B, E), and decreased telomere number (Fig. S7C, F), phenotypes similar to those that resulted from tethering SUMO to telomeres shown in our previous study (Zhao et al., 2024). Again, this effect was not seen with MMS21(C215A) or PIAS4 (C342A) mutants (Fig. S7A-F, Movie 7, 8), confirming the need for SUMOylation for phase separation. In all assays, the two SUMO ligases gave comparable results (Fig. S6B, D, F, H), indicating that SUMOylation on telomeres rather than ligase identity is important for APB formation and other ALT features. Similar observations were made in PML KO U2OS cells (Fig. S6B-D), though SUMO intensity per unit of ligase recruited to telomeres (Fig. S6G) and the degree of telomere clustering were less compared with those seen in PML-containing cells (Fig. S6F). This result indicates that PML helps to enrich SUMO on telomeres to promote telomere clustering.

**Fig. 5.**
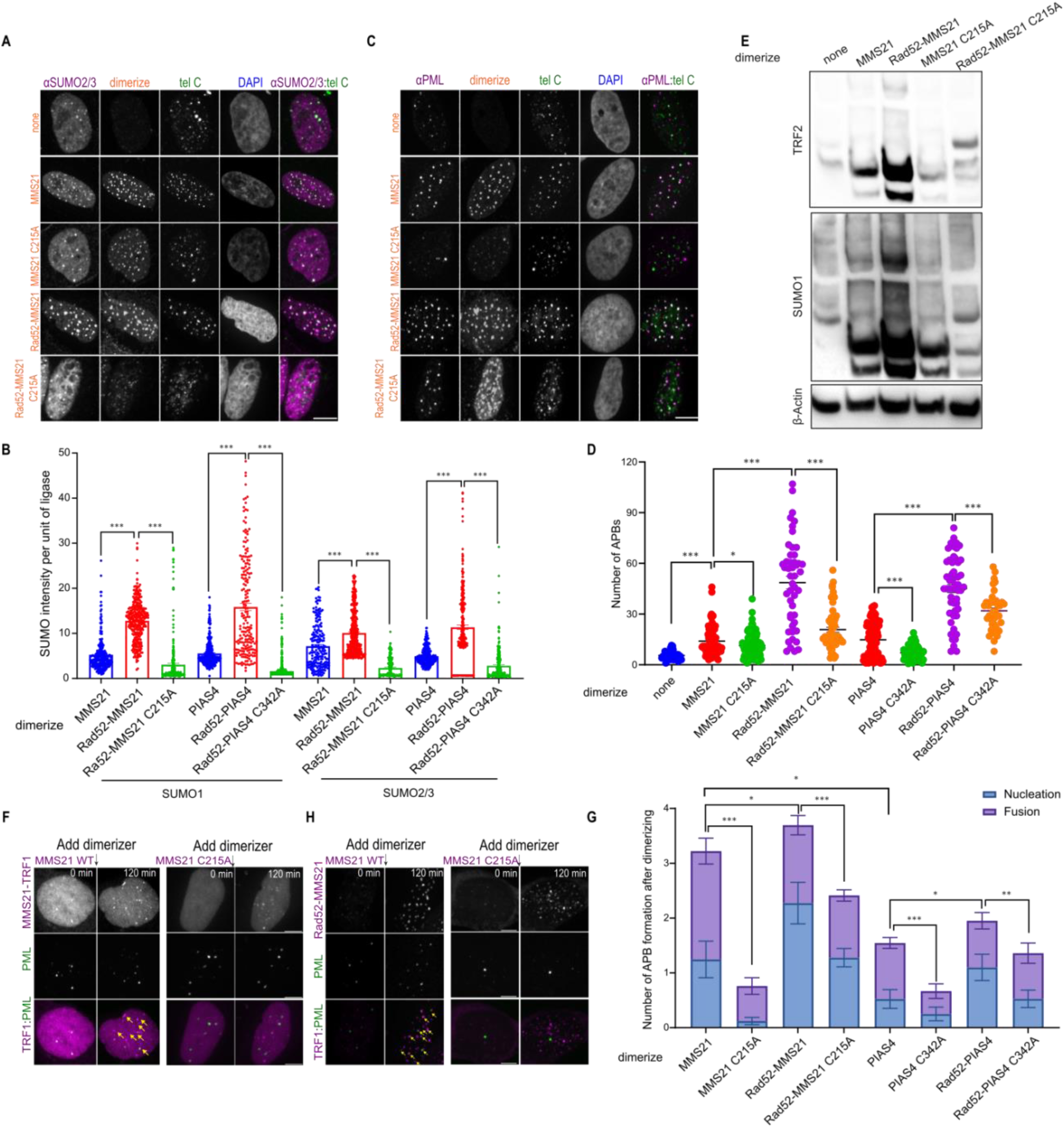
Recruiting SUMO ligases fused to Rad52 enhances APB nucleation. **(A)** Representative images and **(B)** quantification of SUMO1/2/3 intensity per unit of MMS21/PIAS4 or Rad52-MMS21/PIAS4/MMS21(C215A)/PIAS4(C342A) intensity on telomeres in U2OS cells. Each dot represents one cell in three independent experiments, more than 47 cells were analyzed in each group. **(C)** Representative images and **(D)** quantification of PML localization at telomeres in U2OS cells after dimerizing MMS21, MMS21(C215A), Rad52-MMS21/PIAS4 or Rad52-MMS21(C215A)/PIAS4(C342A) for 6 hours. Each dot represents one cell in three independent experiments, more than 55 cells were analyzed in each group. **(E)** U2OS cells were transfected with HA-SUMO3, 3xHalo-GFP-TRF1 and mCherry-eDHFR-MMS21/MMS21(C215A)/Rad52-MMS21/Rad52-MMS21(C215A). Cells were immunoblotted with anti-TRF2, SUMO1 and β-Actin. **(F)** Representative images of cells with endogenous PML labeled with Clover, expressing mCh-eDHFR-MMS21/MMS21(C215A) or **(H)** Rad52-MMS21/MMS21(C215A) and 3xHalo-TRF1 with the addition of the dimerizer at indicated time points. Yellow arrows indicate APBs. **(G)** Quantification of newly formed APBs per cell after dimerizing MMS21(C215A)/Rad52-MMS21/PIAS4(C342A)/Rad52-PIAS4(C342A) for 2 hours. More than 25 cells were analyzed in each group from three independent experiments. Scale bars, 5 μm.

Finally, using PML-Clover containing cells to follow APB formation after ligase tethering, we found that MMS21 and PIAS4 targeting to telomeres led to APB formation both by fusion and nucleation (Fig. 5F, G. Fig. S7G, H), while MMS21(C215A) or PIAS4(C342A) had minimal effects (Fig. 5F, G. Fig. S7G, H), confirming the observation from fixed imaging that SUMOylation is necessary for APB formation (Fig. 5C, D). MMS21’s effects were slightly more potent than PIAS4 during the 2-hour of live imaging (Fig. 5G), which differs from the fixed imaging after 5 hours of dimerization that showed a similar level of APB formation (Fig. 5D).

This difference may indicate that MMS21 is more efficient at promoting APB formation at short time scales, likely caused by differences in ligase activity and/or available substrates. Moreover, recruiting both ligases led to APB formation by fusion more than by nucleation (Fig. 5F, G, Fig. S7G, H), similar to the outcomes upon TRF1-FokI induction. The total number of APBs formed was also similar to that induced by TRF1-FokI (Fig. 1G), indicating that SUMO enrichment by ligase recruitment is comparable with telomere break induction. Together, these data show that the enrichment of the two SUMO ligases is equally sufficient to induce APB formation but with fusion more so than nucleation.

## Recruiting Rad52 together with SUMO ligases increases APB nucleation

The combined results using different SUMO enrichment or SUMO depletion regimes are consistent among themselves and support the notion that telomeres with lower SUMO levels support APB fusion but that a higher SUMO level is required for APB nucleation. If this is true, one would predict that while APB nucleation is less prominent than APB fusion in the SUMO ligase telomere targeting regime, further increased telomere-SUMO levels could result in more APB nucleation. To test this prediction, we targeted ligases together with Rad52, a repair protein implicated in ALT that can increase ALT telomere SUMO levels, as shown previously (Zhao et al., 2024). We confirmed that TRF2 SUMOylation increased upon dual tethering of MMS21 and Rad52 (Rad52-MMS21) to telomeres compared to only MMS21, by co-expressing HA-SUMO3 (Fig. 5E). Consistently, SUMO1 and SUMO2/3 intensities per unit of MMS21 and PIAS4 recruited were higher when Rad52 was introduced compared to when just MMS21 and PIAS4 or their enzymatically dead variants MMS21(C215A) and PIAS4(C342A) were targeted to telomeres (Fig. 5A, B, Fig. S6A). Significantly, dual targeting Rad52 and ligases to telomeres was associated with more APB formation (Fig. 5C, D). However, regarding the degree of telomere clustering, the dual targeting system has a comparable effect as ligase targeting alone (Fig. S6H), likely because the SUMO level is no longer important for telomere clustering once the threshold for phase separation, which is required for telomere clustering, is reached. Finally, compared with MMS21 or PIAS4 alone, dual targeting of Rad52 with ligases led to more and faster APB nucleation than recruiting MMS21 or PIAS4 alone (Fig. 5G, H, Fig. S7I-K). This result supports the prediction from our model that higher SUMO levels favor APB formation via nucleation (Fig. 1G).

### APB nucleation is associated with stronger BLM recruitment to telomeres and ALT features

Finally, we address possible functional consequences of the two APB formation pathways. Given the established role of APBs in recruiting BLM to ALT telomeres (Loe et al., 2020), we examine BLM enrichment on telomeres in different situations. As shown above, telomere recruitment of SUMO3, Rad52-MMS21, and Rad52-PIAS4 mainly leads to APB nucleation. For simplicity, we will refer to these tethering strategies as APB nucleation regimes. On the other hand, recruiting SIM, MMS21, and PIAS4 to telomeres mainly results in APB fusion and, thus, is collectively referred to as APB fusion regimes. Immunofluorescence showed that APB nucleation regimes led to more BLM on telomeres than the APB fusion regimes (Fig. 6A, B). This was confirmed using an APEX proximity labeling assay with an APEX-Halo-TRF1 stable U2OS cell line to examine BLM associated with biotinylated TRF1 (Fig. 6D). In addition, APB nucleation regimes led to more telomeric DNA synthesis than the APB fusion regimes (Fig. 6A, C, Fig. S8A). Collectively, these results indicate that APB nucleation is more efficient at localizing BLM to telomeres for ALT telomeric DNA synthesis (Fig. 6E top).

**Fig. 6.**
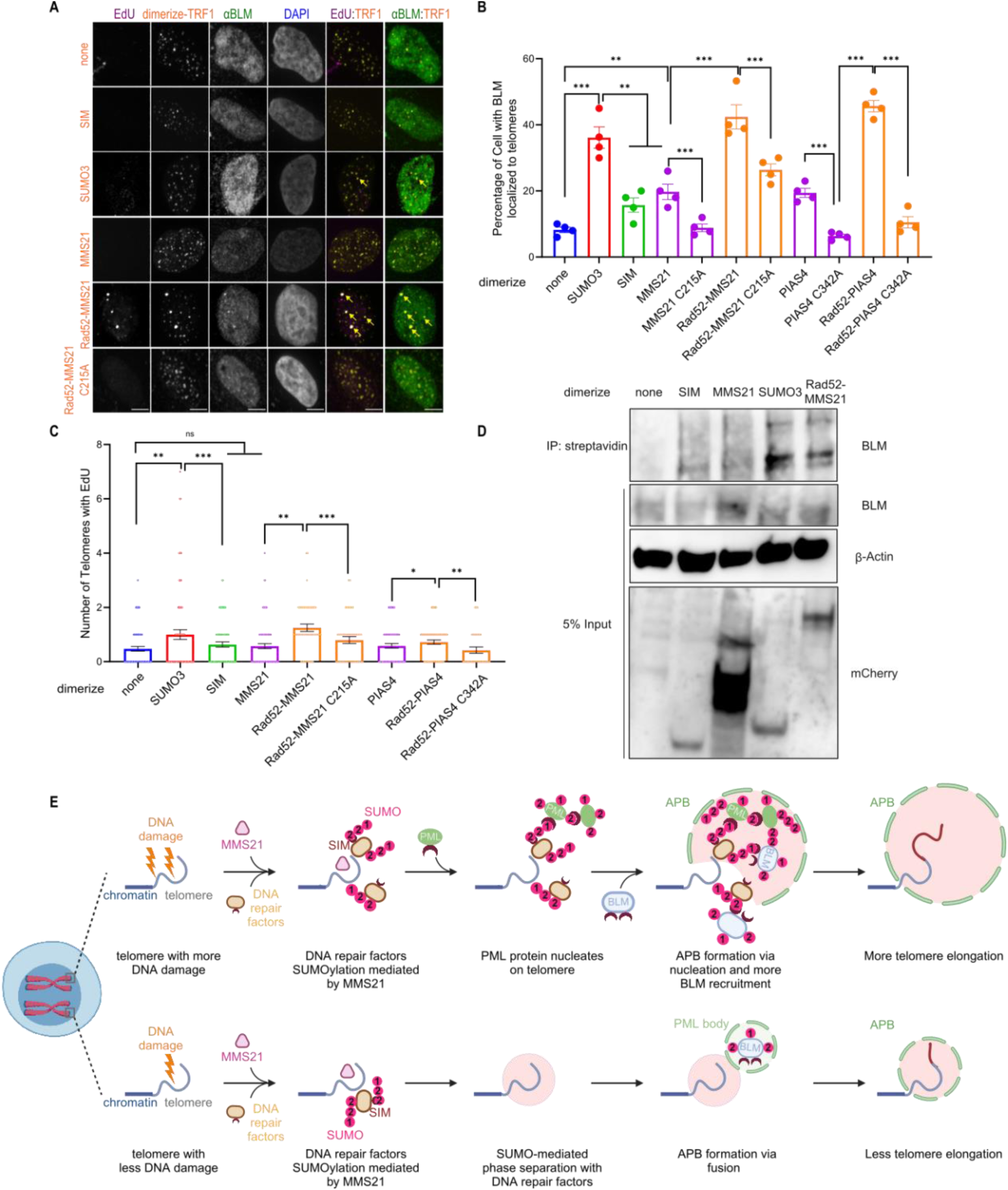
APB nucleation localizes BLM to telomeres more efficiently. **(A)** Representative images of EdU, BLM staining on telomeres after dimerizing SUMO3/SIM/MMS21/Rad52-MMS21 in U2OS cells for 6 hours. Yellow errors indicate EdU signal on telomeres or BLM localization on telomeres. Scale bars, 5 μm. **(B)** Quantification of endogenous BLM localization on telomeres after dimerizing SUMO3/SIM/MMS21/Rad52-MMS21/PIAS4/Rad52-PIAS4 in U2OS cells for 6 hours. Each dot represents one independent experiment, more than 24 cells were analyzed in each group for three independent experiments. **(C)** Quantification of EdU staining on telomeres after dimerizing SUMO3, SIM, MMS21, PIAS4, Rad52-MMS21, Rad52-MMS21(C215A), Rad52-PIAS4, or Rad52-PIAS4(C342A) for 6 hours. Each dot represents one cell in three independent experiments, and more than 51 cells were analyzed in each group. **(D)** APEX-Halo-TRF1 U2OS cells were transfected with mCherry-eDHFR-SIM/MMS21/SUMO3/Rad52-MMS21. Biotinylated telomere proteins were captured with streptavidin beads. Cells were immunoblotted with anti-BLM, mCherry and β-Actin. **(E)** Proposed model. **Top:** Telomeres with more DNA damage recruit more DNA repair factors as SUMO substrates, resulting in the enrichment of SUMO on telomeres by MMS21-mediated polySUMOylation, which then recruits PML protein to nucleate PML NBs on telomeres to promote efficient BLM recruitment and telomere elongation. **Bottom:** Telomeres with less DNA damage recruit less DNA repair factors as SUMO substrates, resulting in low enrichment of SUMO on telomeres, which cannot recruit PML protein to nucleate PML NBs but can fuse with PML NBs to form APBs, resulting in less efficient BLM recruitment and telomere elongation. Created with BioRender.com.

To examine whether the key outcome of the APB nucleation regime is to enrich BLM on telomeres, we asked whether overexpressing BLM could rescue the lower level of telomeric DNA synthesis in the APB fusion regimes. This was indeed the case, as we found that more telomeres were enriched with BLM (Fig. S8B-D) and had more robust telomeric DNA synthesis in the APB fusion regimes after BLM overexpression (Fig. S8E, F). Importantly, these effects were not seen after recruiting MMS21(C215A), PIAS4(C342A), SUMO3(FAAA), SIM(VADA), MMS21(C215A) or PIAS4(C342A), supporting that both SUMO-SIM interaction and SUMOylation are required for BLM mediated telomeric DNA synthesis. Interestingly, BLM overexpression after recruiting MMS21 and PIAS4 resulted in both a similar increase in BLM localization on telomeres (Fig. S8D) and DNA synthesis (Fig. S8F) as recruiting SUMO3, indicating that overexpressing BLM is sufficient to compensate for the defects of BLM recruitment and telomeric DNA synthesis due to the lack of APB nucleation.

## Discussion

Previous studies have revealed the requirement of SUMOylation in APB formation through modifying the telomere-binding proteins, DNA repair factors, and the PML protein (Brouwer et al., 2009; Chung et al., 2011; Potts & Yu, 2007; H. Zhang et al., 2020). In this study, we followed APB formation dynamics by initiating chemically inducible DNA damage or SUMO tethering to telomeres. Through time-lapse imaging, we reveal two pathways for APB formation: de novo nucleation of PML NBs on telomeres and fusion of telomeres with existing PML NBs (Fig. 6E). Through mutagenesis experiments, we show that both pathways require SUMO-SIM interactions at both the telomere and on the PML protein, highlighting the importance of SUMO-SIM interactions in APB formation.

We also show that both APB formation pathways can occur in the same cell, indicating that the determinant for pathway choice is at the telomere level. Through tethering experiments, we show that the identities of SUMO paralogs and SUMO ligases do not affect APB formation pathway choice. Instead, the SUMO level of telomeres matters: telomeres with high SUMO levels favor APB formation via nucleation, while telomeres with low SUMO levels form APBs via fusion. This is likely because greater SUMO levels on telomeres are needed to compete with the existing PML NBs for the free PML protein in the nucleoplasm for de novo nucleation of PML NBs on telomeres. Supporting the importance of SUMO level, knocking down the polySUMO chain forming SUMO2/3 reduces APB formation via nucleation more than the non-polySUMO chain forming SUMO1. In addition, tethering DNA repair factor Rad52 with SUMO ligases can increase SUMO levels and promote APB formation via nucleation, indicating that the amount of DNA repair factors–likely caused by different degrees of DNA damage on each telomere–can contribute to the pathway choice of APB formation (Fig. 6E). These results agree with the previously reported promiscuous nature of SUMOylation where SUMO ligases modify a group of substrates in the vicinity, and it is this group modification, not the modification of individual proteins, that is functionally important (Dhingra & Zhao, 2019; Hu & Parvin, 2014; Psakhye & Jentsch, 2012; Jentsch & Psakhye, 2013).

We further demonstrate that the functional relevance of the APB formation pathway lies in how it contributes to BLM recruitment to telomeres, one of the key roles of APBs in ALT (Loe et al., 2020). Our data show that both pathways of APB formation can help recruit BLM to telomeres, but APB formation via nucleation has a stronger effect (Fig. 6E). This is likely because telomeres with SUMO levels high enough to compete with the existing PML NBs for free PML protein are also likely to form bigger PML NBs, which in turn recruits more BLM. As a result of more BLM recruitment, telomeres that can form APBs via nucleation have more robust telomeric DNA synthesis. We suspect that telomeres with APBs formed via nucleation are likely to elongate more than telomeres with APBs formed via fusion in the same cell, leading to heterogeneity in telomere length that is often observed in ALT cells. In supporting this prediction, ALT cells that lack APBs are shown to lose telomere length heterogeneity (Loe et al., 2020).

Our findings may explain why PML NBs are associated with damaged ALT telomeres more often than non-telomere DNA damage sites that also have SUMO accumulation (Chang et al., 2018; Dhingra & Zhao, 2019; Hu & Parvin, 2014; Psakhye & Jentsch, 2012; Yeung et al., 2012). First, ALT telomeres are known to experience high replication stress that can result in DNA damage (Lu & Pickett, 2022). This can lead to greater enrichment of repair proteins that can be SUMOylated at telomeres. In addition, ALT cells have some extremely long telomeres (Bryan et al., 1995). These long telomeres harbor more telomere-binding proteins that can be promiscuously SUMOylated in response to damage-induced SUMO ligase localization to telomeres. Through either SUMOylation of repair proteins or telomere binding proteins, the enhanced SUMO accumulation can promote PML NB nucleation on telomeres. On the other hand, telomeres are relatively mobile compared to other regions of the genome, which offers a high chance for the SUMO-coated telomeres to collide with PML NBs. Lastly, with its repetitive DNA and heterochromatin nature, telomeres might be difficult to repair, leading to persistent damage foci with a higher opportunity to accumulate SUMO and/or fuse with PML NBs. In supporting this kinetic model for PML NB-damage site association, PML NBs are not found at easy-to-repair DNA or early stage of damage sites, but a stable association of PML NBs with persistently damaged foci can be observed at difficult-to-repair damage sites (Hornofova et al., 2023; Vancurova et al., 2019; Rodier et al., 2011; Kleijwegt et al., 2023).

In summary, our work fills several gaps in our knowledge regarding APB biogenesis in ALT. The results may also conceptually impact models of the interplay between SUMO and PML NBs in other pathways, such as DNA replication (Stubbe et al., 2020) and transcription (Pavan Kumar et al., 2007; Tan et al., 2008; Gasser and Stutz. 2023; Salsman et al., 2017), where both SUMO and PML NBs are known to play regulatory roles. In addition, our discoveries on the mechanisms of PML NB localization to ALT telomeres may contribute to the understanding of other condensate-chromatin associations, such as the dynamic contacts between nuclear speckles and transcriptional sites (Zhong et al., 2020).

## Materials and Methods

### Cell culture

All experiments were performed with U2OS cells unless otherwise stated. PML KO U2OS cells were gifts from Dr. Eros Lazzerini Denchi. SUMO1 KO, SUMO2 KO and SUMO2 KO with SUMO2 rescue cells were gifts from Dr. Michael J Matunis. U2OS cells with endogenous PML tagged with Clover (U2OS Clover-PML) were previously described (Pinder et al., 2015). Cells were cultured in growth medium (Dulbecco’s Modified Eagle’s medium with 10% FBS and 1% penicillin–streptomycin) at 37 °C in a humidified atmosphere with 5% CO2.

### Plasmids

The plasmid for inducing DNA damage at telomeres (mCherry-TRF1-FokI) was previously published (Cho et al., 2014). 3xHalo-GFP-TRF1 was previously published (H. Zhang et al., 2020). SUMO1/2/3 (or SUMO mutant) for mCherry-eDHFR-SUMO came from plasmids gifted by Karsten Rippe (Chung et al., 2011). SIM and SUMO mutants for mCherry-eDHFR-SIM/SUMO were from plasmids gifted by Michael Rosen, where the SIM interacting mutants of SUMO were generated by mutating the FKIK (SUMO3) or FKVK (SUMO1) to FAAA, non-conjugatable SUMO modules were generated by mutating the C-terminal di-Gly motif to di-Val, and the mutant of SIM defective in SUMO binding was generated by mutating the VIDL sequence into VADA (Banani et al., 2016). The vector containing mCherry-eDHFR was from our published plasmid Mad1-mCherry-eDHFR (H. Zhang et al., 2017). PML WT, PML SUMOylation sites were from plasmids gifted by Michael Rosen (Banani et al., 2016). PML SIM mutant was kindly provided by Dr. Hung-Ying Kao and subcloned into the target backbone. GFP-BLM was from Addgene plasmid #80070. GFP-MMS21, PIAS4 and mutants were introduced into target plasmids through in-fusion cloning (#638948, Takara Bio). HA-SUMO3 plasmid was a gift from Dr. Roderick O’Sullivan. All the target plasmids in this study were derived from a plasmid that contains a CAG promoter for constitutive expression, obtained from E. V. Makeyev (Khandelia et al., 2011).

### Synchronize cells to G2

Cells were first treated with 2 mM thymidine (T1895, Sigma-Aldrich) for 21 h, released into fresh medium for 4 h, and then treated with 15 μM CDK1i (RO-3306, cat# SML0569, Sigma-Aldrich) for 12 h.

### siRNA transfection

siRNA for SUMO1 (sc-29498, Santa cruz), SUMO2/3 (sc-37167, Santa cruz), MMS21 (L-018070-00-0005, Horizon), and PIAS4 (L-006445-00-0005, Horizon) were purchased commercially. Cells were transfected with 100 nM of siRNA and RNAiMax (13778075, ThermoFisher) diluted in OptiMEM (31-985-070, Life Technologies). The transfection medium was replaced with culture media 8 hr later and the transfection was repeated on day 2. Cells were collected and imaged at 48 hr after the second-round transfection.

### Protein dimerization and FokI induction

Design, synthesis, and storage of the dimerizer TMP (trimethoprim)-Fluorobenzamide-Halo ligand (TFH) were previously reported (Lackner et al., 2022). Dimerization to telomeres was performed as previously described (Zhao et al., 2021). Briefly, TFH was added directly to the growth medium to a final working concentration of 100 nM. Cells were incubated with the dimerizer-containing medium for the indicated times, followed by immunofluorescence (IF) or fluorescence in situ hybridization (FISH). For live imaging with protein dimerization, the dimerizers were first diluted to 200 nM in growth medium and then added to cell chambers to the working concentration after first-round imaging.

To induce damage on telomere in cells transfected with mCherry-ER-DD-TRF1-FokI, Shield-1 (Cheminpharma LLC), and 4-hydroxytamoxifen (4-OHT) (Sigma-Aldrich) at 1 μM were added for 2h to allow TRF1 to enter the nucleus after first-round imaging, as previously described (Cho et al., 2014).

### Immunofluorescence (IF) and telomere DNA fluorescence in situ Hybridization (FISH)

IF and FISH were performed following a previously published protocol (Zhao et al., 2021). Cells were fixed in 4% formaldehyde for 10 min at room temperature, followed by permeabilization in 0.5% Triton X-100 for 10 min. Cells were incubated with primary antibody at 4°C in a humidified chamber overnight and then with secondary antibody for 1 h at room temperature before washing and mounting. Primary antibodies were anti-SUMO1 (Ab32058, Abcam,1:200 dilution), anti-SUMO2/3 (Asm23, Cytoskeleton, 1:200 dilution), anti-PML (sc966, Santa Cruz, 1:100 dilution), anti-BLM (A300-110A, Bethyl, 1:100 dilution), anti-PCNA (2586S, Cell signaling, 1:1000 dilution), and anti-RPA (NB600-565, Novus, 1:100 dilution). For IF-FISH, coverslips were first stained with primary and secondary antibodies, then fixed again in 4% formaldehyde for 10 min at room temperature. Coverslips were then dehydrated in an ethanol series (70%, 80%, 90%, 2 min each) and incubated with a TelC-Alexa488 or TelC-Cy3 PNA probe (F1004, F1002, Panagene) at 75°C for 5 min and then overnight in a humidified chamber at room temperature. Coverslips were then washed and mounted for imaging.

### Telomeric DNA synthesis detection by EdU

Following transfection, cells were pulsed with EdU (10 μM) along with protein dimerization or DNA damage induction for 6 hrs before harvest. Cells on glass coverslips were washed twice in PBS and fixed with 4% paraformaldehyde (PFA) for 10 min. Cells were permeabilized with 0.3% (v/v) Triton X-100 for 5 min. The Click-IT Plus EdU Cell Proliferation Kit with Alexa Flour 647 (C10635, Invitrogen) was applied to cells for 30 minutes to detect EdU.

### Cell imaging and image processing

Imaging acquisition was performed as previously described (Xu et al., 2024). For live imaging, cells were seeded on 22 x 22 mm glass coverslips coated with poly-D-lysine (P1024, Sigma-Aldrich). When ready for imaging, coverslips were mounted in magnetic chambers (Chamlide CM-S22-1, LCI) with cells maintained in a normal medium supplemented with 10% FBS and 1% penicillin/streptomycin at 37 °C on a heated stage in an environmental chamber (TOKAI HIT Co., Ltd.). Images were acquired with a microscope (ECLIPSE Ti2) with a 100 × 1.4 NA objective, a 16 XY Piezo-Z stage (Nikon Instruments Inc.), a spinning disk (Yokogawa), an electron multiplier charge-coupled device camera (IXON-L-897), and a laser merge module that was equipped with 488 nm, 561 nm, 594 nm, and 630 nm lasers controlled by NIS-Elements Advanced Research. For both the fixed cells and live imaging, images were taken with 0.5 μm spacing between Z slices, for a total of 8 μm. For movies, images were taken at 5 min intervals for up to 3 hr.

Images were processed and analyzed using NIS-Elements AR (Nikon). Maximum projections were created from z stacks, and thresholds were applied to the resulting 2D images to segment and identify telomere/SUMO foci as binaries. For quantification of the co-localization of two fluorescent labels, images were analyzed using binary operations in NIS-Elements AR. Co-localized foci were counted if the objects from different layers contained overlapping pixels.

### Western blotting

Cells were harvested with trypsin, quickly washed in PBS, and directly lysed in 4XNuPage LDS sample buffer (NP0007, Thermo). The resulting whole-cell lysates were analyzed by Western blotting with the following primary antibodies: SUMO1 (Ab32058, Abcam,1:1000 dilution), SUMO2/3 (Asm23, Cytoskeleton, 1:1000 dilution), MMS21 (A304-129A, Bethyl, 1:1000 dilution), PIAS4 (4392, Cell Signaling, 1:1000 dilution), TRF2 (NBP1-86911, Novus, 1:1000 dilution), mCherry (ab183628, Abcam, 1:1000 dilution), BLM (A300-110A, Bethyl, 1:1000 dilution), HA (3724S, Cell signaling, 1:1000 dilution) and β-Actin (Ab8226, Abcam, 1:1000 dilution). The HRP signal was visualized with Super Signal ECL substrate (34095, Pierce) per the manufacturer’s instructions.

### SUMOylation protein pull-down

Approximately 4×10^6^ cells were scraped and lysed with 300 μL NP40 buffer (FNN0021, Thermo) with protease inhibitor (78430, Thermo) and N-Ethylmaleimide (NEM) (E3876, Sigma) to prevent SUMO deconjugation. A fraction (20 μL) of the cell suspensions was kept as input. 3 uL of HA antibody (3724S, Cell signaling) was added to each group and was rotated at 4°C for 2 hours. Protein A/G beads (88802, Thermo) (12 μL for each sample) were washed three times with NP40 buffer and then were incubated together with the lysates and antibody at 4°C overnight. Subsequently, Protein A/G beads were washed three times. Finally, the proteins captured by Protein A/G beads were eluted by boiling in LDS Sample Buffer (NP0007, Invitrogen). The levels of indicated proteins in input and pull-down samples were analyzed by western blot.

### Proximity Biotinylation Using the Peroxidase APEX2

Biotin phenol (SML2135, Sigma) was dissolved in dimethyl sulfoxide as a 500 mM stock solution and was added directly to cell culture media to a final concentration of 500 μM. H_2_O_2_ (H1009, Sigma Aldrich) was spiked into the cell culture media to a final concentration of 1 mM to induce biotinylation. After 1 min of very gentle swirling, the media was decanted as quickly as possible and the cells were washed three times with phosphate-buffered saline (PBS) containing 100 mM sodium azide (S2002, Sigma), 100 mM sodium ascorbate (PHR1279, Sigma), and 50 mM TROLOX (6-hydroxy-2,5,7,8-tetramethylchroman-2-carboxylic acid) (AC218940010, Tocris). Approximately 4×10^6^ cells were scraped and lysed with 300 μL RIPA buffer (89900, Thermo) with protease inhibitor (78430, Thermo) and N-Ethylmaleimide (NEM) (E3876, Sigma) to prevent SUMO deconjugation. A fraction (20 μL) of the cell suspensions was kept as input. Streptavidin C1 beads (65001, Invitrogen) (12 μL for each sample) were washed three times with RIPA denaturing buffer and were then incubated together with the lysates at 4°C overnight. Subsequently, Streptavidin C1 beads were washed three times. Finally, the proteins captured by Streptavidin C1 beads were eluted by boiling in LDS Sample Buffer (NP0007, Invitrogen). The levels of indicated proteins in input and pull-down samples were analyzed by western blot.

### Statistical methods

All error bars represent means ± SEM. Statistical analyses were performed using Prism 10.0 (GraphPad software). Two-tailed unpaired t-tests have been used for all tests. Statistical significance: N.S., not significant, P > 0.05; *P < 0.05; **P < 0.01; ***P < 0.001.

## Acknowledgments

We thank Dr. Roger A. Greenberg (University of Pennsylvania), Dr. Eros Lazzerini Denchi (National Cancer Institute), Dr. Michael Rosen (UT Southwestern), Dr. Karsten Rippe (German Cancer Research Center), Dr. Roderick J. O’Sullivan (University of Pittsburgh) and Dr. Hung-Ying Kao (Case Western Reserve University) for kindly gifting plasmids and cell lines. This work was supported by the United States National Institutes of Health Grant U01CA260851 to HZ, GM118510 to DC, R35 GM145260 to XZ., GM060980 to MM, a Project Grant from the Canadian Institutes of Health Research (CIHR) PJT-156017 to GD.

## Author contributions

H.Z., R.Z. conceptualized this study. R.Z. designed and conducted the experiments. R. L. and D.C. designed and synthesized the dimerizers. J. S. and G. D. made the U2OS PML-Clover cell line. R.Z., A.W., and H.Z., analyzed the results. R.Z. made the figures. R.Z., and H.Z. wrote the manuscript with comments from other authors.

## Competing interests

The authors declare no competing interests.

## Data availability

All data needed to evaluate the conclusions in the paper are present in the paper and the Supplementary Materials.

## Supplementary Materials

**Movie 1** Inducing DNA damage at telomeres in PML-Clover cells. Movie for Fig. 1C. Left: Composite of FokI-TRF1 (magenta) and PML (green), middle: mCh-FokI-TRF1, right: PML. 4-OHT and shield 1 were added to cells after the first time point to induce damage. The boxes show APBs formed by fusion and nucleation. Scale bars, 5 μm.

**Movie 2** Recruiting SUMO3 to telomeres in PML-Clover cells. 3xHalo-TRF1 was co-transfected. Movie for Fig. 1I. Left: Composite of SUMO3 (magenta) and PML (green), middle: mCh-eDHFR-SUMO3, right: PML. The dimerizer was added to cells after the first time point to induce dimerization. The box shows APBs formed by fusion and nucleation. Scale bars, 5 μm.

**Movie 3** Recruiting SUMO3(FAAA) to telomeres in PML-Clover cells. 3xHalo-TRF1 was co-transfected. Movie for Fig. S1C. Left: Composite of SUMO3(FAAA) (magenta) and PML (green), middle: mCh-eDHFR-SUMO3(FAAA), right: PML. The dimerizer was added to cells after the first time point to induce dimerization. Scale bars, 5 μm.

**Movie 4** Recruiting SIM to telomeres in PML-Clover cells. 3xHalo-TRF1 was co-transfected. Movie for Fig. S1D. Left: Composite of SIM (magenta) and PML (green), middle: mCh-eDHFR-SIM, right: PML. The dimerizer was added to cells after the first time point to induce dimerization. The box shows an APB formed by fusion. Scale bars, 5 μm.

**Movie 5** Dimerizing MMS21 to telomeres in U2OS cells. Movie for Fig. S7A. Left: Composite of MMS21 (magenta) and TRF1 (green), middle: mCh-eDHFR-MMS21, right: 3xHalo-GFP-TRF1. The dimerizer was added to cells after the first time point to induce dimerization. The box shows a telomere fusion event. Scale bars, 5 μm.

**Movie 6** Dimerizing MMS21(C215A) to telomeres in U2OS cells. Movie for Fig. S7A. Left: Composite of MMS21(C215A) (magenta) and TRF1 (green), middle: mCh-eDHFR-MMS21(C215A), right: 3xHalo-GFP-TRF1. The dimerizer was added to cells after the first time point to induce dimerization. Scale bars, 5 μm.

**Movie 7** Dimerizing PIAS4 to telomeres in U2OS cells. Movie for Fig. S7D. Left: Composite of PIAS4 (magenta) and TRF1 (green), middle: mCh-eDHFR-PIAS4, right: 3xHalo-GFP-TRF1. The dimerizer was added to cells after the first time point to induce dimerization. The box shows a telomere fusion event. Scale bars, 5 μm.

**Movie 8** Dimerizing PIAS4(C342A) to telomeres in U2OS cells. Movie for Fig. S7D. Left: Composite of PIAS4(C342A) (magenta) and TRF1 (green), middle: mCh-eDHFR-PIAS4(C342A), right: 3xHalo-GFP-TRF1. The dimerizer was added to cells after the first time point to induce dimerization. Scale bars, 5 μm.

**Fig. S1.**
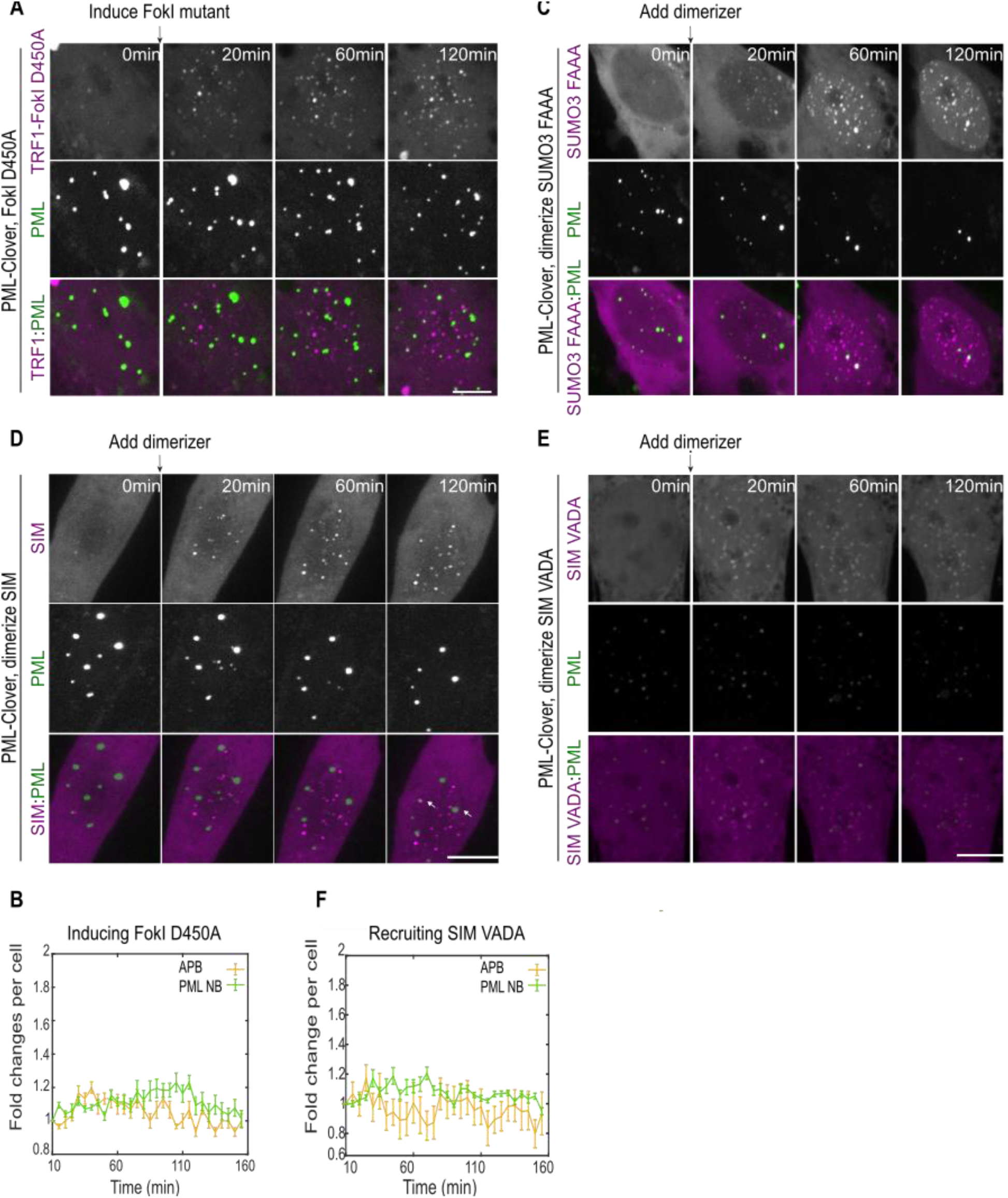
Both pathways of APB formation require SUMO-SIM interaction. **(A)** Representative images of cells with endogenous PML labeled with Clover and expressing mCh-TRF1-FokI D450A with the treatment of 4-Hydroxyestradiol (4-OHT) and shield 1 to induce FokI at indicated time points. **(B)** Quantification of APB and PML NB numbers per cell after FokI D450A induction (numbers are normalized by the number at the first time point for each cell, more than 17 cells per group, three independent experiments). **(C)** Representative images of cells with endogenous PML labeled with Clover, expressing mCh-eDHFR-SUMO3(FAAA) and 3xHalo-TRF1 with adding the dimerizer at indicated time points. **(D)** Representative images of cells with endogenous PML labeled with Clover, expressing mCh-eDHFR-SIM and 3xHalo-TRF1 with adding the dimerizer at indicated time points. **(E)** Representative images of cells with endogenous PML labeled with Clover, expressing mCh-eDHFR-SIM(VADA) and 3xHalo-TRF1 with adding the dimerizer at indicated time points. **(F)** Quantification of APB and PML NB numbers per cell after adding the dimerizer when recruiting SIM(VADA) to PML-Clover telomeres (numbers are normalized by the number at the first time point for each cell, more than 14 cells per group, three independent experiments). Scale bars, 5 μm.

**Fig. S2.**
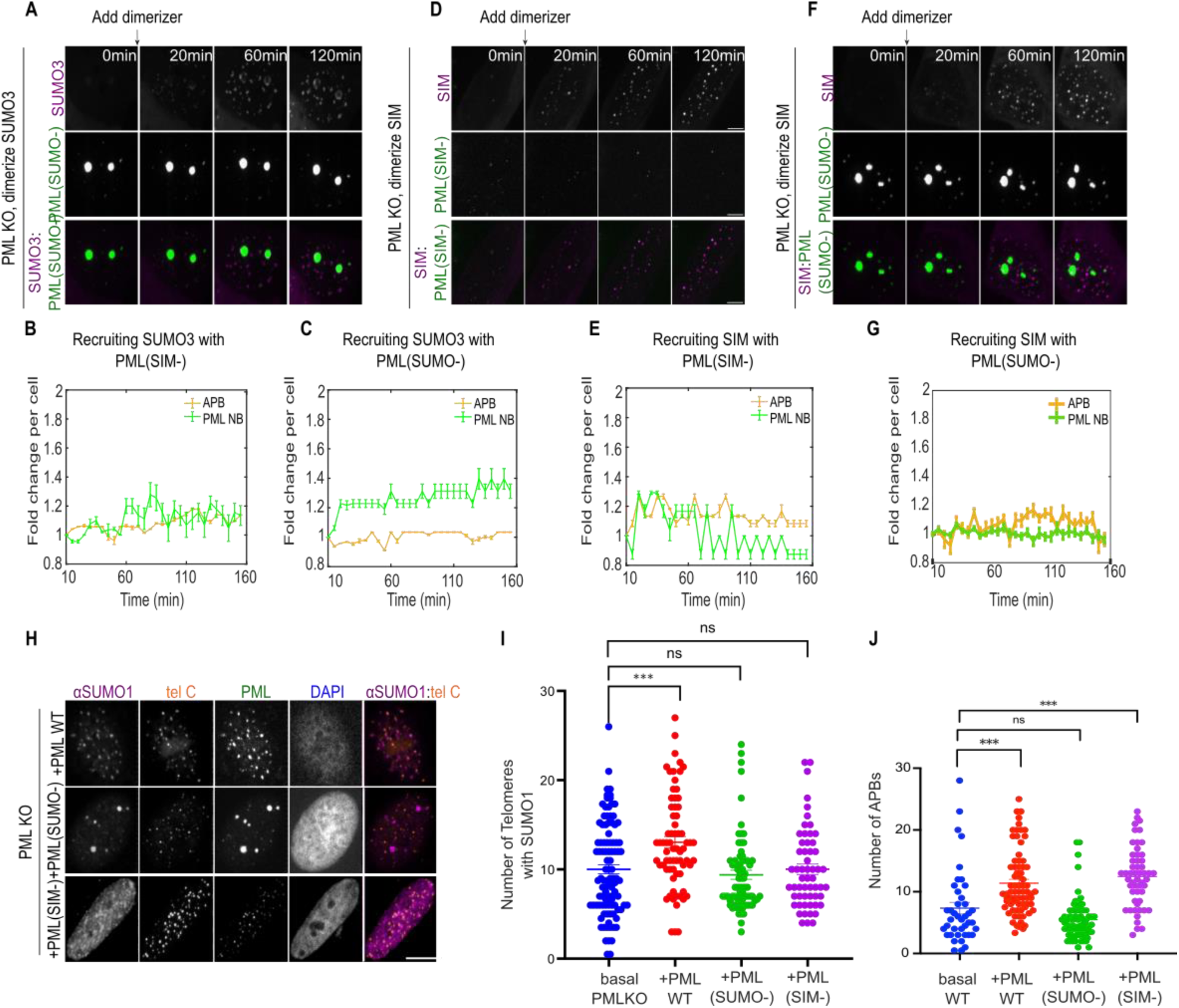
PML contributes to ALT via its SUMOylation sites and SIM. **(A)** Representative images of PML KO cells expressing mCh-eDHFR-SUMO3, 3xHalo-TRF1, and PML(SUMO-) with the dimerizer to dimerize at indicated time points. **(B)** Quantification of APB and PML NB numbers per cell after adding the dimerizer when recruiting SUMO3 in PML KO cells expressing PML(SIM-) and **(C)** PML(SUMO-). **(D)** Representative images of PML KO cells expressing mCh-eDHFR-SIM, 3xHalo-TRF1, and PML(SIM-) with the dimerizer to dimerize at indicated time points. **(E)** Quantification of APB and PML NB numbers per cell after adding the dimerizer when recruiting SIM in PML KO cells expressing PML(SIM-) and **(G)** PML(SUMO-). Numbers in (B)(C)(E)(G) are normalized by the number at the first time point for each cell, more than 17 cells per group, three independent experiments. **(F)** Representative images of PML KO cells expressing mCh-eDHFR-SIM, 3xHalo-TRF1, and PML(SUMO-) with the dimerizer to dimerize at indicated time points. **(H)** Representative images and **(I)** quantification of SUMO1 localization on telomeres in G2 arrested PML KO after overexpressing PML WT, PML(SUMO-) or PML(SIM-). Each dot represents one cell in three independent experiments, more than 54 cells were analyzed in each group. **(J)** Number of APBs in G2 arrested PML KO after overexpressing PML WT, PML(SUMO-), or PML(SIM-). Each dot represents one cell in three independent experiments, more than 45 cells were analyzed in each group. Scale bars, 5 μm.

**Fig. S3.**
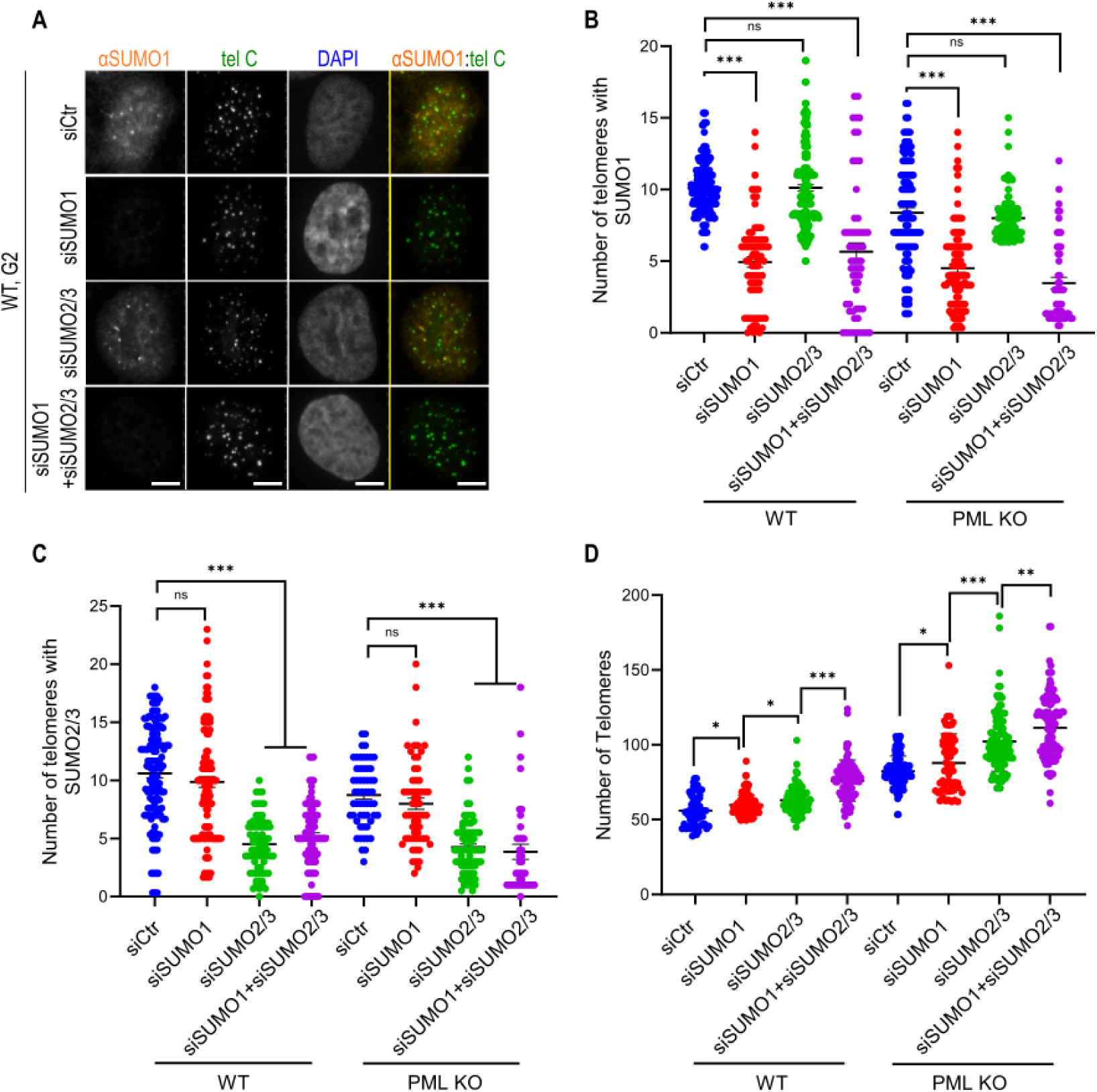
siRNA knocking down reveals that SUMO2/3 contributes more to ALT phenotypes than SUMO1. **(A)** Representative images and **(B)** quantification of SUMO1 localization on telomeres in G2 arrested U2OS WT after transfection of control siRNA, siSUMO1 or siSUMO2/3 for two days. Each dot represents one cell in three independent experiments, more than 56 cells were analyzed in each group. **(C)** Quantification of SUMO2/3 localization on telomeres in G2 arrested U2OS WT after transfection of control siRNA, siSUMO1 or siSUMO2/3 for two days. Each dot represents one cell in three independent experiments, more than 50 cells were analyzed in each group. **(D)** Quantification of telomere number in G2 arrested U2OS WT after transfection of control siRNA, siSUMO1 or siSUMO2/3 for two days. Each dot represents one cell in three independent experiments, more than 56 cells were analyzed in each group. Scale bars, 5 μm.

**Fig. S4.**
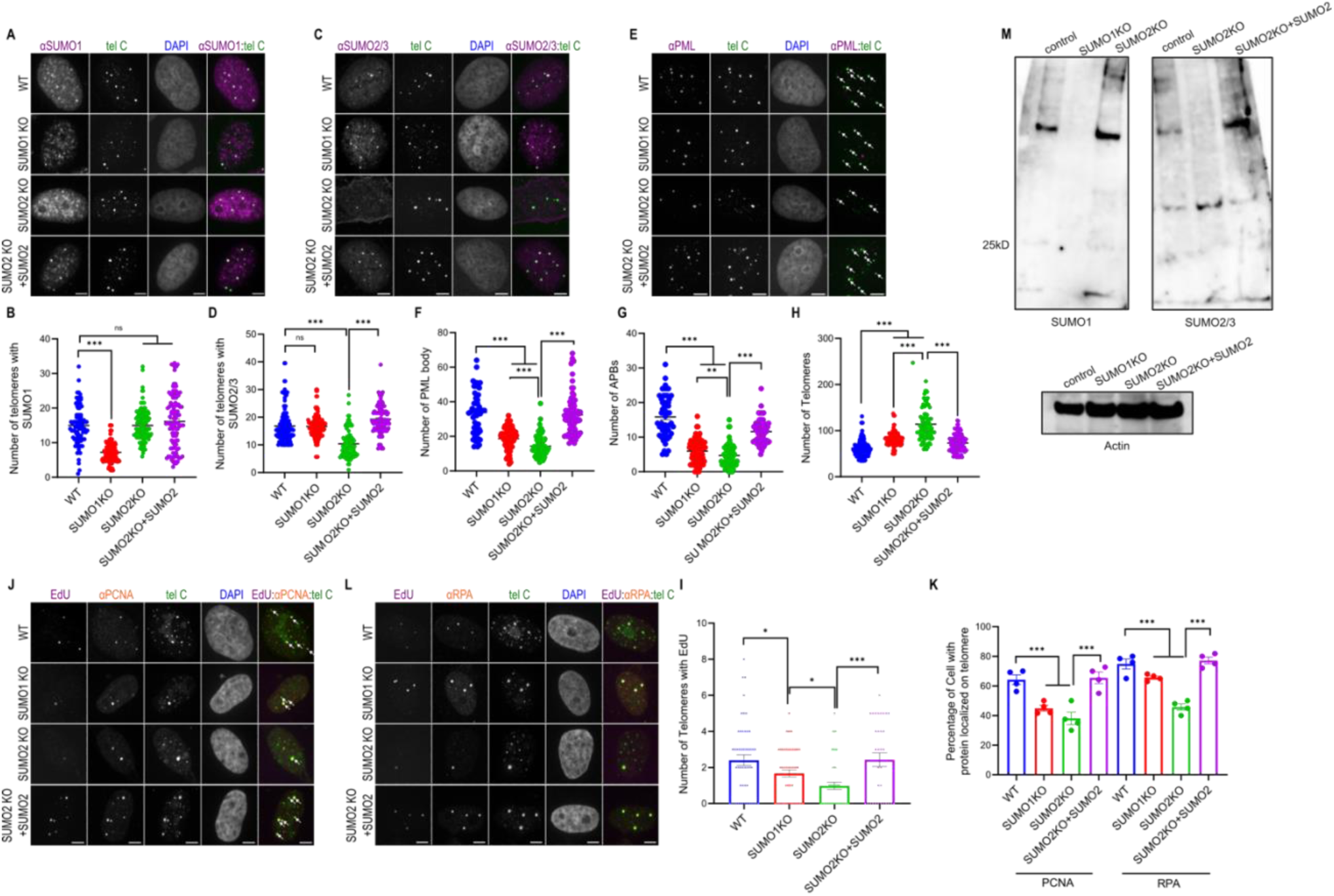
SUMO knockout reveals that SUMO2/3 contributes more to ALT phenotypes than SUMO1. **(A)** Representative images and **(B)** quantification of SUMO1 localization on telomeres in G2 arrested U2OS WT, SUMO1 KO, SUMO2 KO, and SUMO2 KO+SUMO2 cells. Each dot represents one cell in three independent experiments, more than 52 cells were analyzed in each group. **(C)** Representative images and **(D)** quantification of SUMO2/3 localization on telomeres in G2 arrested U2OS WT, SUMO1 KO, SUMO2 KO, and SUMO2 KO+SUMO2 cells. Each dot represents one cell in three independent experiments, more than 55 cells were analyzed in each group. **(E)** Representative images and **(F)** quantification of PML localization on telomeres in G2 arrested U2OS WT, SUMO1 KO, SUMO2 KO, and SUMO2 KO+SUMO2 cells. Each dot represents one cell in three independent experiments, more than 50 cells were analyzed in each group. **(G)** Quantification of total PML NBs in G2 arrested U2OS WT, SUMO1 KO, SUMO2 KO, and SUMO2 KO+SUMO2 cells. Each dot represents one cell in three independent experiments, more than 50 cells were analyzed in each group. **(H)** Quantification of telomere number in G2 arrested U2OS WT, SUMO1 KO, SUMO2 KO, and SUMO2 KO+SUMO2 cells. Each dot represents one cell in three independent experiments, more than 53 cells were analyzed in each group. **(I)** Quantification of EdU on telomeres in G2 arrested U2OS WT, SUMO1 KO, SUMO2 KO, and SUMO2 KO+SUMO2 cells. Each dot represents one cell in three independent experiments, more than 42 cells were analyzed in each group. **(J)** Representative images of EdU and PCNA or **(L)** RPA on telomeres in G2 arrested U2OS WT, SUMO1 KO, SUMO2 KO, and SUMO2 KO+SUMO2 cells. **(K)** Quantification of endogenous PCNA and RPA localization on telomeres in G2 arrested U2OS WT, SUMO1 KO, SUMO2 KO, and SUMO2 KO+SUMO2 cells. Each dot represents one independent experiment, more than 22 cells were analyzed in each group for three independent experiments. **(M)** Western blot of SUMO1, SUMO2/3 or β-Actin in SUMO1 KO, SUMO2 KO, SUMO2 KO+SUMO2, and control U2OS cells. Scale bars, 5 μm.

**Fig. S5.**
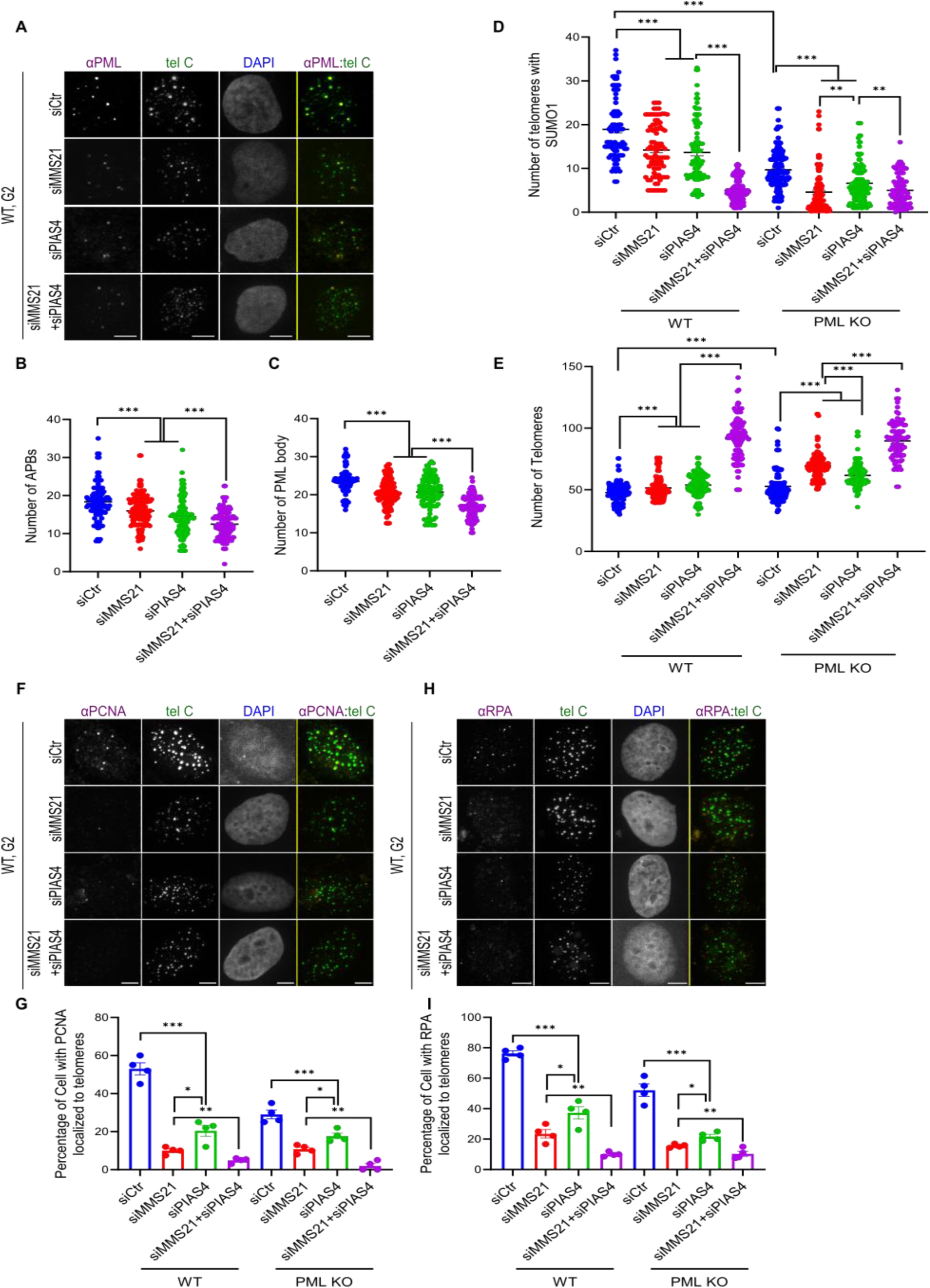
MMS21 contributes more to SUMO on telomeres than PIAS4 in the absence of PML. **(A)** Representative images and **(B)** quantification of PML localization on telomeres in G2 arrested U2OS WT after transfection of control siRNA, siMMS21, or siPIAS4 for two days. Each dot represents one cell in three independent experiments, more than 45 cells were analyzed in each group. **(C)** Quantification of total PML NBs in G2 arrested U2OS WT after transfection of control siRNA, siMMS21, or siPIAS4 for two days. Each dot represents one cell in three independent experiments, more than 46 cells were analyzed in each group. **(D)** Quantification of SUMO1 localization on telomeres and **(E)** telomere number in G2 arrested U2OS WT and PML KO cells after transfection of control siRNA, siMMS21, or siPIAS4 for two days. Each dot represents one cell in three independent experiments, more than 53 cells were analyzed in each group. **(F)** Representative images and **(G)** quantification of PCNA localization on telomeres in G2 arrested U2OS WT after transfection of control siRNA, siMMS21, or siPIAS4 for two days. Each dot represents one independent experiment, more than 23 cells were analyzed in each group for three independent experiments. **(H)** Representative images and **(I)** quantification of RPA localization on telomeres in G2 arrested U2OS WT after transfection of control siRNA, siMMS21, or siPIAS4 for two days. Each dot represents one independent experiment, more than 28 cells were analyzed in each group for three independent experiments. Scale bars, 5 μm.

**Fig. S6.**
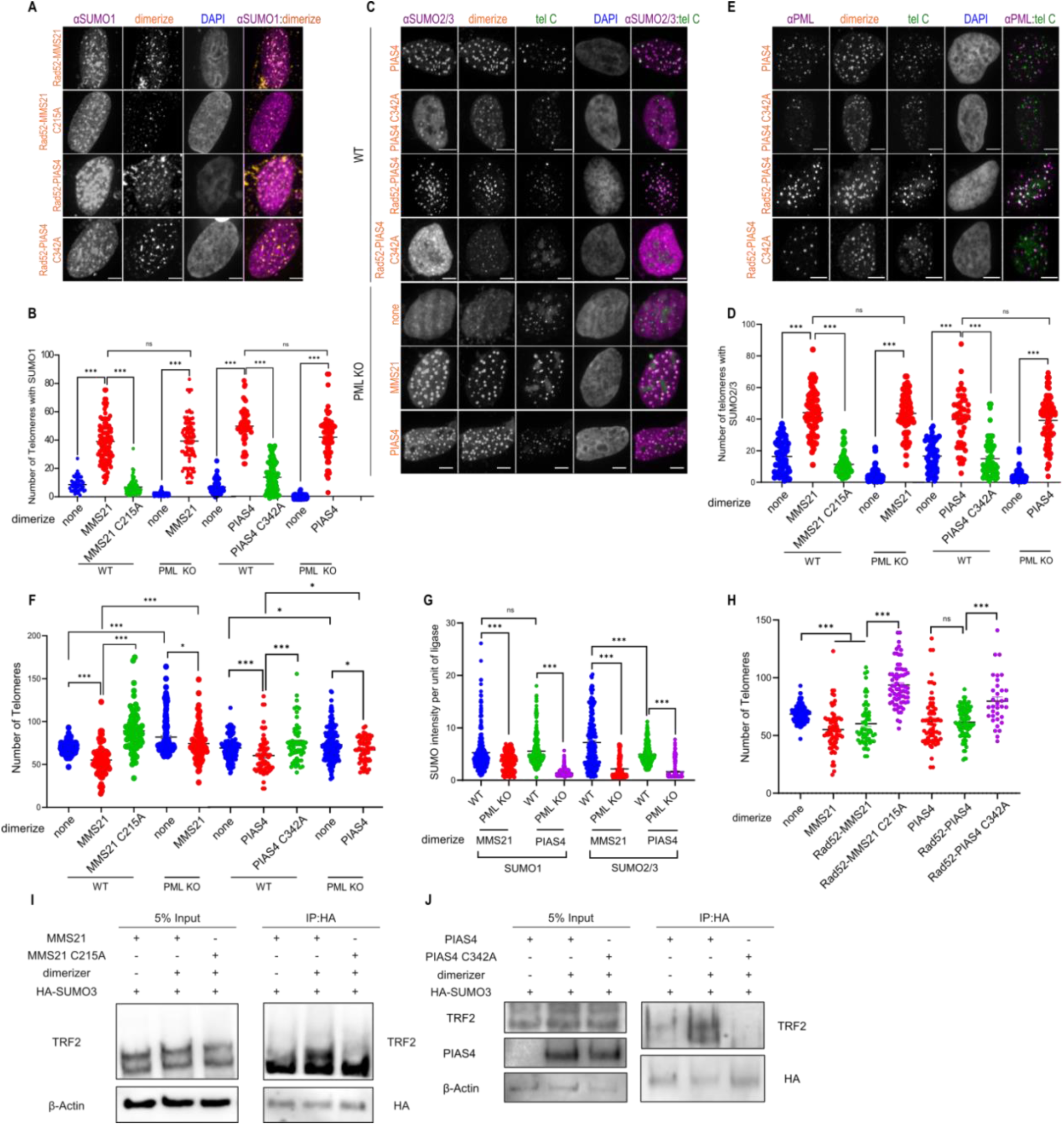
Recruiting SUMO ligases is sufficient for APB formation. **(A)** Representative images and **(B)** quantification of SUMO1 localization at telomeres in U2OS WT and PML KO cells after dimerizing MMS21(C215A)/PIAS4(C215A) to telomeres for 6 hours. Each dot represents one cell in three independent experiments, more than 54 cells were analyzed in each group. **(C)** Representative images and **(D)** quantification of SUMO2/3 localization at telomeres in U2OS WT and PML KO cells after dimerizing MMS21 or PIAS4 to telomeres for 6 hours. Each dot represents one cell in three independent experiments, more than 54 cells were analyzed in each group. **(E)** Representative images of PML localization on telomeres in U2OS WT after dimerizing PIAS4, PIAS4(C342A), Rad52-PIAS4, or Rad52-PIAS4(C342A) to telomeres for 6 hours. **(F)** Quantification of telomere number in U2OS WT and PML KO cells after dimerizing MMS21, PIAS4, MMS21(C215A), or PIAS4(C215A) to telomeres for 6 hours. Each dot represents one cell in three independent experiments, more than 55 cells were analyzed in each group. **(G)** Quantification of SUMO1/2/3 intensity per unit of MMS21 or PIAS4 intensity on telomeres in U2OS WT and PML KO cells. Each dot represents one cell in three independent experiments, more than 52 cells were analyzed in each group. **(H)** Quantification of telomere number in U2OS WT cells after dimerizing MMS21/PIAS4/Rad52-MMS21(C215A)/Rad52-PIAS4(C342A) to telomeres for 6 hours. Each dot represents one cell in three independent experiments, more than 46 cells were analyzed in each group. **(I)** U2OS cells expressing Halo-GFP-TRF1, HA-SUMO3, mCherry-eDHFR-MMS21(C215A), or **(J)** mCherry-eDHFR-PIAS4(C342A) with adding 5-hour of dimerizer. SUMOylated proteins were captured with protein A/G beads following HA antibody incubation. Levels of TRF2 and HA in inputs and HA pull-downs were analyzed by western blot. Scale bars, 5 μm.

**Fig. S7.**
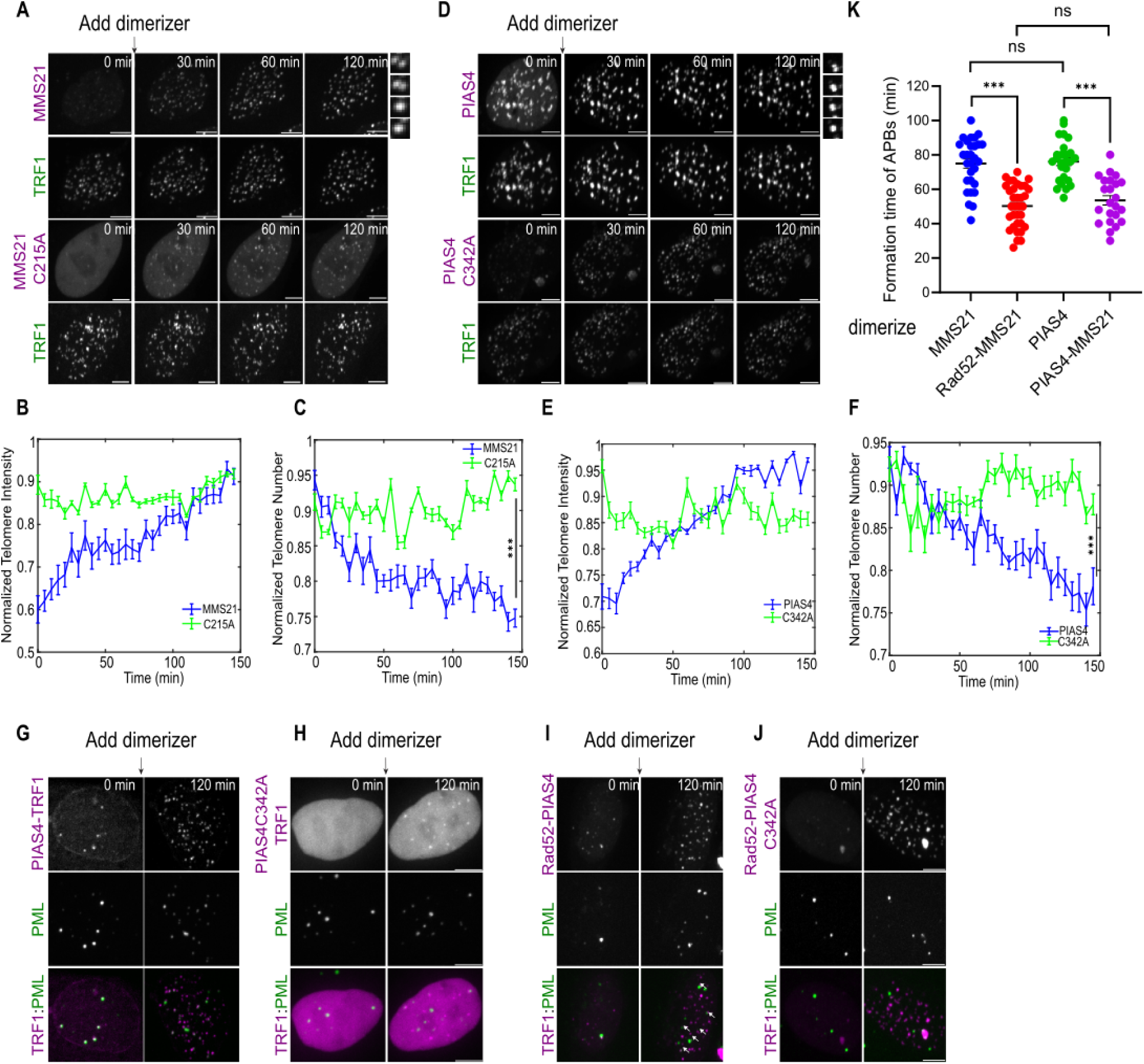
Recruiting SUMO ligases is sufficient for phase separation on telomeres. **(A)** Representative images of U2OS cells after dimerizing mCh-eDHFR-MMS21 or MMS21(C215A) to 3xHalo-GFP-TRF1 after the first time point. Zoomed-in images on the right show the fusion of TRF1 foci after dimerizing MMS21. **(B)** Telomere sum intensity and **(C)** telomere number per cell after dimerizing MMS21 or MMS21(C215A) to U2OS telomeres (more than 21 cells per group, three independent experiments, two-tailed unpaired t-test). **(D)** Representative images of U2OS cells after dimerizing mCh-eDHFR-PIAS4 or PIAS4(C342A) to 3xHalo-GFP-TRF1 after the first time point. Zoomed-in images on the right show the fusion of TRF1 foci after dimerizing PIAS4. **(E)** Telomere sum intensity and **(F)** Telomere number per cell after dimerizing PIAS4 or PIAS4(C342A) to U2OS telomeres (more than 19 cells per group, three independent experiments, two-tailed unpaired t-test). **(G)** Representative images of cells with endogenous PML labeled with Clover, expressing mCh-eDHFR-PIAS4 or **(H)** mCh-eDHFR-PIAS4(C342A) and 3xHalo-TRF1 with adding the dimerizer at indicated time points. **(I)** Representative images of cells with endogenous PML labeled with Clover, expressing mCh-eDHFR-Rad52-PIAS4 or **(J)** mCh-eDHFR-Rad52-PIAS4(C342A) and 3xHalo-TRF1 with adding the dimerizer at indicated time points. **(K)** APB formation time after dimerizing MMS21/Rad52-MMS21/PIAS4/Rad52-PIAS4 for 2 hours. Each dot represents one cell in three independent experiments, more than 30 cells were analyzed in each group. Scale bars, 5 μm or 1 μm for cropped images.

**Fig. S8.**
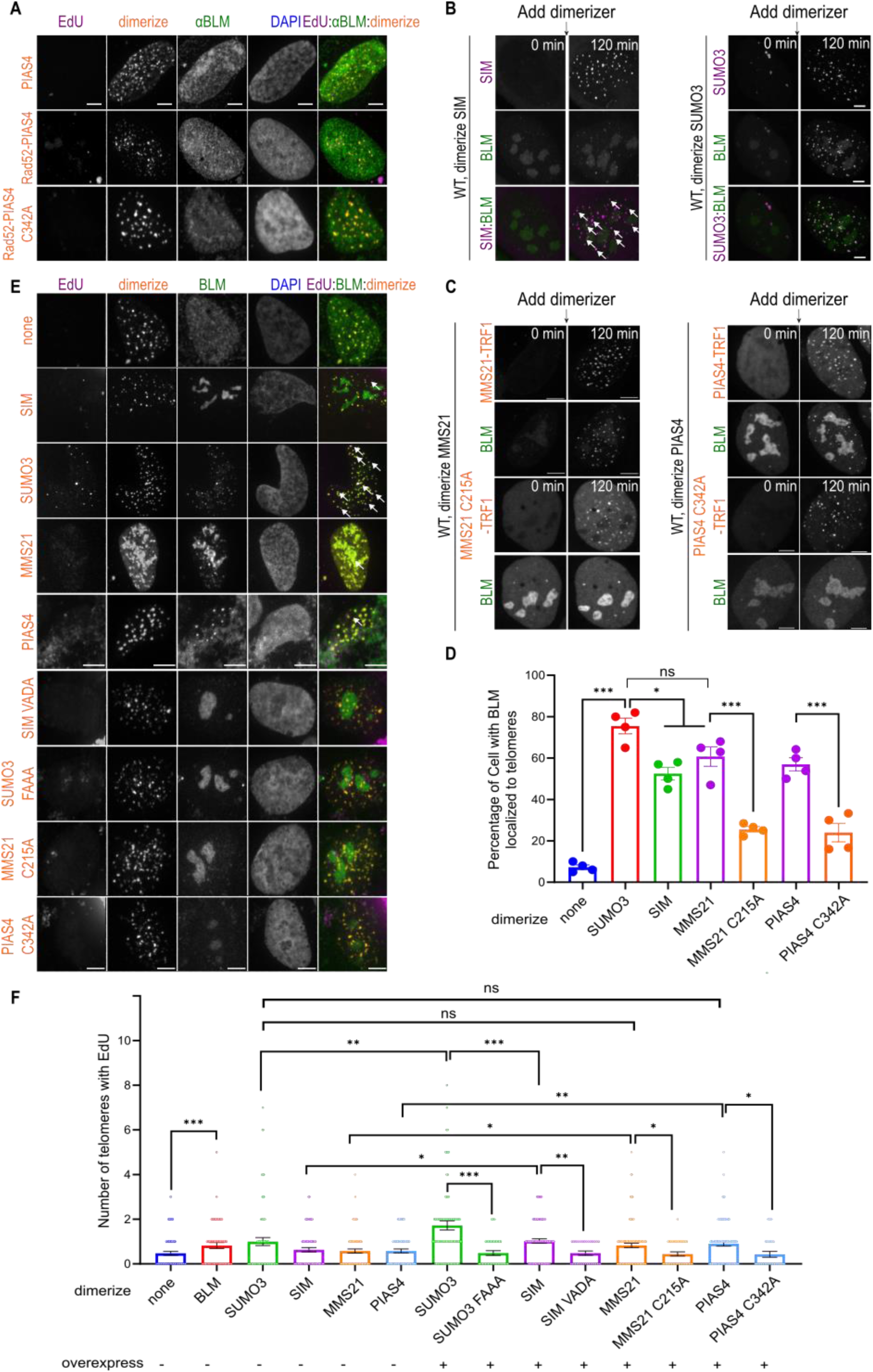
Recruitment of Rad52 together with ligases promotes nucleation of APB and telomeric DNA synthesis. **(A)** Representative images of EdU, BLM staining on telomeres after dimerizing PIAS4/Rad52-PIAS4/Rad52-PIAS4(C342A) in U2OS WT cells for 6 hours. **(B)** Representative images of BLM localization at telomeres after dimerizing mCh-eDHFR-SIM or SUMO3 to 3xHalo-TRF1 in U2OS cells expressing GFP-BLM at indicated time. **(C)** Representative images of BLM localization at telomeres after dimerizing mCh-eDHFR-MMS21/MMS21(C215A) or PIAS4/PIAS4(C342A) to 3xHalo-TRF1 in U2OS cells expressing GFP-BLM at indicated time. **(D)** Quantification of overexpressed BLM localization on telomeres after dimerizing SUMO3/SIM/MMS21(C215A)/PIAS4(C342A) to telomeres in U2OS cells for 6 hours. Each dot represents one independent experiment, more than 28 cells were analyzed in each group for three independent experiments. **(E)** Representative images of EdU staining on telomeres after dimerizing SUMO3/SIM/MMS21/PIAS4 or SUMO3(FAAA)/SIM(VADA)/MMS21(C215A)/PIAS4(C342A) for 6 hours, with GFP-BLM overexpression. Yellow arrows indicate EdU localization telomeres. **(F)** Quantification of EdU staining on telomeres after dimerizing SUMO3/SIM/MMS21/PIAS4 or SUMO3(FAAA)/SIM(VADA)/MMS21(C215A)/PIAS4(C342A) to telomeres for 6 hours, with GFP-BLM overexpression. Each dot represents one cell in three independent experiments, more than 47 cells were analyzed in each group. Scale bars, 5 μm.

## Notes

### Competing Interest Statement

The authors have declared no competing interest.

